# Sodium coordination at the Na2 site enforces ion-substrate coupling in serotonin transporter SERT

**DOI:** 10.64898/2026.11.19.767907

**Authors:** Eva Hellsberg, Danila Boytsov, Jasper D. Shide, Eray Esendir, Annika Backer, Jiahui Huang, Nathan Bernhardt, Walter Sandtner, Lucy R. Forrest

**Affiliations:** National Institutes of Neurological Disorders and Stroke, National Institutes of Health; Bethesda MD 20892 USA; Center of Physiology and Pharmacology, Department of Pharmacology, Medical University of Vienna; Vienna 1090 Austria

## Abstract

Secondary active transporters use the thermodynamically favorable movement of an ion down its electrochemical gradient to drive substrate accumulation against a concentration gradient. For the human serotonin transporter (SERT), strict coupling between Na⁺ and 5-HT⁺ transport is essential for maintaining neurotransmitter homeostasis, yet the structural mechanism by which Na⁺ binding enforces this coupling is incompletely understood. Here, we identify an interaction network through which Na⁺ coordination at the cation site Ct2 (also known as Na2) modulates conformational change on both sides of the transporter. Molecular dynamics simulations reveal that Na⁺ occupancy of Ct2 stabilizes specific interactions between residues in TM5, TM8, and TM1 that together gate aqueous access to the cytoplasmic and extracellular pathways. Disruption of this network through mutations at Ct2 residues Asp437 and Ser438, or at the TM5 residue Thr284, reduces the concentrative power of SERT and allows a constitutive Na⁺ conductance, consistent with uncoupled Na⁺ uniport. These findings reveal how structural integrity at and around the Ct2 site enforces obligatory Na⁺–5-HT⁺ coupling and thereby sustains the thermodynamic efficiency of serotonin uptake.

## Introduction

Among the integral membrane proteins that act as secondary active transporters, the monoamine transporters, including serotonin (5-hydroxytryptamine, 5-HT^+^) transporter SERT (SLC6A4), are among the best studied. The monoamine transporters form a branch of the SLC6 family that also includes transporters for dopamine (DAT/SLC6A3), and norepinephrine (NET/SLC6A2). SLC6 transporters use symport of Na^+^ to harvest the electrochemical Na^+^ gradient as a driving force for substrate translocation across the cellular membrane (Kristensen et al. 2011). SERT gains an additional driving force for 5-HT^+^ uptake by coupling to antiport of K^+^ under physiological conditions, or H^+^ *in vitro* (Nelson and Rudnick 1979; Rudnick and Nelson 1978; Keyes and Rudnick 1982; Hasenhuetl, Freissmuth, and Sandtner 2016; Hellsberg et al. 2024). In addition, SERT requires extracellular Cl^−^ to achieve transport (Lingjaerde 1971; Schicker et al. 2012; Huang et al. 2026). Notably, recent work suggests that, in addition to their well-known impacts on mood, SERT inhibitors support anti-tumor immunity (Li et al. 2025), and impact cardiovascular (Shen et al. 2025) and gastrointestinal health (Acharekar et al. 2022), emphasizing the need to understand SERT function at its most basic level.

All secondary active transporters operate according to the alternating access mechanism. Accordingly, substrate can bind to the transporter from one or the other side of the lipid membrane but not from both sides simultaneously. Thus, the protein must be able to adopt at least two major conformational states in which the central pocket is accessible for substrate binding: an outward- and an inward-open conformation. Conformational changes between these states (**Fig. 1A**) obey specific rules that ensure net transport of substrate occurs under physiological conditions. Due to the high concentrations of Na^+^ and Cl^−^ in the extracellular milieu, in the absence of serotonin, SERT favors an extracellular-open conformation bound to at least one Na^+^ ion and one Cl^−^ ion (Fenollar-Ferrer et al. 2014; Coleman, Green, and Gouaux 2016). When available, 5-HT^+^ binds to the exposed central substrate binding site, known as S1 (Yang and Gouaux 2021). Once 5-HT^+^ is bound, the transporter is able to convert first to an occluded conformation (Gradisch et al. 2022; Chan et al. 2022) and then to an inward-open state (Zhang and Rudnick 2006; Schicker et al. 2012; Yang and Gouaux 2021; Chan et al. 2022). In the latter, the binding sites are exposed to the cytoplasm (Androutsellis-Theotokis, Ghassemi, and Rudnick 2001; Forrest et al. 2008; Zhang et al. 2018) where the concentration of Na^+^ is low, favoring Na^+^ dissociation. In the absence of bound Na^+^, the affinity of the inward-open transporter for serotonin is also reduced and therefore substrate unbinds (Koldso et al. 2011; Chan et al. 2022) leading to accumulation. Lastly, a K^+^ ion, which is at a high concentration internally, binds to the inward-open conformation (Rudnick and Nelson 1978; Nelson and Rudnick 1979; Hellsberg et al. 2024; Zhao and Tajkhorshid 2026) and reduces the energetic barrier of conversion to the outward-open state (Hasenhuetl, Freissmuth, and Sandtner 2016). Once the substrate sites are again exposed to the extracellular milieu, where the K^+^ concentration is low, K^+^ is expected to unbind. *In vitro*, if internal K^+^ is omitted, protons take on the role of accelerating the return step (Keyes and Rudnick 1982; Hasenhuetl, Freissmuth, and Sandtner 2016; Hellsberg et al. 2024). Although extracellular Cl^−^ binding is required (Lingjaerde 1971; Tavoulari et al. 2011), Cl^−^ gradients cannot drive substrate transport (Huang et al. 2026). The complete transport cycle, and its physiological rates and efficiency, are therefore largely determined by the concentration gradients of the three cations Na^+^, K^+^, and H^+^, relative to that of 5-HT^+^.

**Figure 1.**
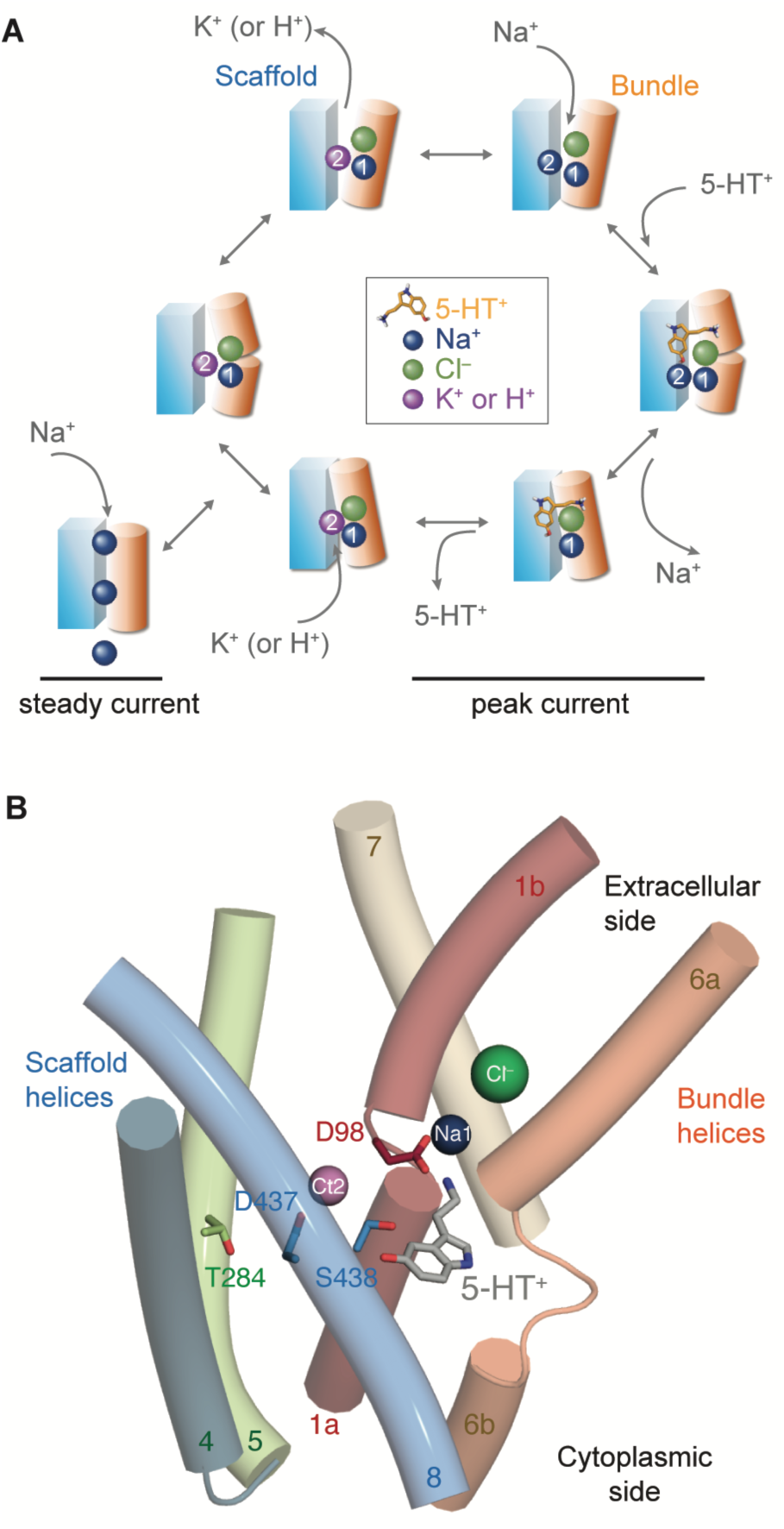
Transport cycle and structure of human serotonin transporter, SERT. (**A**) The protein cycles between outward-facing (at top), occluded and inward-facing conformations, largely involving movements of helices in the bundle (*orange cylinder*) against helices in the scaffold (*blue cuboid*). We posit that Na^+^ ion (*blue sphere*, at the Na1 site, “1”) and one Cl^−^ ion (*green sphere*) are bound in all transitions of the cycle. Due to its high concentration on the external side, a second Na^+^ ion will bind to the Ct2 site (“2”) which favors conformations with the cytoplasmic pathway closed and the extracellular pathway open. When available, 5-HT^+^ (*sticks*) can bind to the S1 site (including in low Na^+^ concentrations). Occupation of both Ct2 and S1 sites favors closure of the extracellular pathway. Opening of the cytoplasmic pathway is associated with release of one Na^+^, followed by 5-HT^+^, which is associated with the inward peak current observed electrophysiologically. The “return” steps that reset the protein to an outward-facing state are rate-limiting and accelerated by a cation, typically K^+^ (or H^+^ at low pH), *purple sphere*. When K^+^ (or H^+^ at low pH) binds to the inward-facing state, SERT can access an off-cycle, channel-like state (*lower left*) that supports steady Na^+^ conductance. (**B**) Key elements of the SERT outward-open structure (Protein Databank identifier 5I6X). Selected helices from bundle (TM1, TM6, and TM7; *oranges/reds*) and scaffold (TM4, TM5 and TM8; *greens/blues*) are shown as cylinders. Substrate 5-HT^+^ and side chains for Asp98, Thr284, Asp437 and Ser438 are shown as sticks, with carbon atoms colored by the respective helix, or gray for substrate. Figure made using PyMol v2.1.3 (Schrödinger, Inc.).

SLC6 transporters belong to the broader family of neurotransmitter:sodium symporters (NSS), which adopt the so-called LeuT fold (Yamashita et al. 2005) characterized by twelve membrane-spanning helices that can be considered to comprise bundle and scaffold regions (Forrest and Rudnick 2009; Licht et al. 2024). A major rotation of the bundle transmembrane (TM) helices TM1, TM2, TM6, and TM7 around an axis passing through the substrate sites orchestrates the alternating exposure of the central binding site, together with discrete movements of transmembrane helices TM1 and TM6 and the intracellular loop connecting TM4 to TM5 (Malinauskaite et al. 2014; Coleman, Green, and Gouaux 2016; Yang and Gouaux 2021; Gradisch et al. 2022; Licht et al. 2024). Reported structures of LeuT, DAT, NET, and SERT (Yamashita et al. 2005; Penmatsa, Wang, and Gouaux 2015; Wang, Penmatsa, and Gouaux 2015; Coleman, Green, and Gouaux 2016; Yang and Gouaux 2021; Song and Wu 2024; Srivastava et al. 2024), supported by molecular dynamics (MD) simulations of SERT(Hellsberg et al. 2019; Szollosi and Stockner 2021; Gradisch et al. 2022; Szollosi and Stockner 2022; Chan et al. 2022; Huang et al. 2026), provide compelling evidence for two binding sites for Na^+^ ions, traditionally referred to as the Na1 and Na2 sites.

The Na2 site lies at the bundle-scaffold interface between TM1 and TM8 (**Fig. 1B**), a location that is critical for stabilizing outward-open conformations in SERT (Tavoulari et al. 2016; Möller et al. 2019) and across the NSS family (Khelashvili et al. 2016; Razavi, Khelashvili, and Weinstein 2017; Stolzenberg et al. 2017). In contrast, Na1 is embedded within the bundle helices adjacent to the Cl^−^ ion site (**Fig. 1B**), suggesting a more localized structural role. Consistent with this placement, perturbations of the Na1 site have comparatively subtle effects on global conformational change in SERT (Felts et al. 2014; Tavoulari et al. 2016; Zhang et al. 2018).

The presence of two Na^+^ binding sites is generally consistent with symport stoichiometries of two or more Na^+^ ions per substrate by NSS transporters specific to amino acids such as glycine and γ-amino-butyric acid, GABA (Roux and Supplisson 2000; Radian and Kanner 1983). That is, two Na^+^ ions bind cooperatively with substrate and release together with the substrate but are not bound in the return transition. However, early measurements in platelet plasma membrane vesicles indicated a 1:1 Na^+^:5-HT^+^ stoichiometry for SERT (Talvenheimo et al. 1983; Chang and Lam 1998), raising the question of whether both Na^+^ ion sites also participate in transport of serotonin.

Recent observations offer a resolution to this apparent discrepancy. Cl^−^ gradients cannot drive transport (Huang et al. 2026) and Cl^−^ translocation into the cytoplasm is not required for substrate uptake (Hasenhuetl, Freissmuth, and Sandtner 2016), suggesting that Cl^−^ remains bound during the transport cycle. Because the Na1 and Cl⁻ sites are structurally overlapping, sharing coordinating residues Ser336 and Asn368 and placing the ions <8 Å apart (Coleman and Gouaux 2018; Coleman, Green, and Gouaux 2016), their occupancies are likely to be interdependent. Indeed, Poisson-Boltzmann calculations of LeuT indicating strong electrostatic interactions between charges at these sites (Forrest et al. 2007) while MD simulations show co-dependent occupancy (Tavoulari et al. 2016; Huang et al. 2026). Together, these findings are consistent with a model in which both Na1 and Cl sites remain occupied during the return transition (**Fig. 1A**). Additional support for this model is provided by the observation that Ca²⁺-dependent Na1-site mutants do not co-translocate Ca²⁺ ions (Felts et al. 2014).

A corollary of this “bound NaCl” mechanism is that the Na2 site binds the co-transported Na⁺ ion (**Fig. 1A**) and, moreover, is likely involved in the major conformational transitions that underlie Na⁺–5-HT coupling. In other words, Na2 may be a key determinant of the concentrative power of Na⁺ co-transport by SERT (Felts et al. 2014).

During the return transition, the type of bound cation will differ, as Na^+^ is released from Na2 and instead K^+^ (or H^+^) binding is required. The stoichiometry of 5-HT^+^ to K^+^ or to H^+^ antiport is also 1:1 as measured in platelet plasma membrane vesicles (Nelson and Rudnick 1979; Keyes and Rudnick 1982) and recent data indicates that the Na2 site constitutes the binding site for this antiported K^+^ ion (or H^+^) in SERT (Hellsberg et al. 2024). Thus, the Na2 site can be considered multi-purpose and nonspecific and is referred to hereafter as cation site Ct2 (**Fig. 1B**).

Structures of LeuT and SERT reveal that the Ct2 site is structurally disrupted in fully inward-open states, due to separation of TM1a from TM8 (Krishnamurthy and Gouaux 2012; Coleman et al. 2019; Malinauskaite et al. 2014). However, in the MhsT transporter, unwinding of the cytoplasmic TM4-5 loop also creates an aqueous pathway that reaches the Ct2 site in an inward-occluded state, even without TM1a having moved away (Malinauskaite et al. 2014). This unwinding event and pathway have been implicated in Na^+^ permeation to the cytoplasm in other NSS transporters (Malinauskaite et al. 2014; Stolzenberg et al. 2017). In the case of SERT, however, we lack the molecular details of how coupled binding of Na^+^ and 5-HT^+^ controls the opening of the two pathways. Thus, in the present study, we conducted MD simulations of SERT to map out interactions involving the Ct2 site, which predict an important role for an interaction partner in TM5. Electrophysiological measurements of SERT mutants confirm the predicted role for TM5 and, through characterization of constitutive cation currents in these mutants, reveal the mechanistic underpinnings of ion-substrate coupling in Na^+^-driven cotransport of 5-HT^+^.

## Materials and Methods

### MD simulation starting models and overview

Simulations of outward-occluded, outward-open and inward-open conformations of human SERT were performed as described in (Huang et al. 2026) and (Hellsberg et al. 2024), following the protocol of (Tan et al. 2022) and initiated with: a well-curated outward-occluded model constructed from fragments of an outward-open structure, where the extracellular side of the bundle was repositioned according to the outward-occluded structure of LeuT (Hellsberg et al. 2019); an outward-open structure (Protein Database, PDB identifier 5I71 (Coleman, Green, and Gouaux 2016)); or an inward-open structure (PDB identifier 6DZZ (Coleman et al. 2019) excluding residues 78-85 (Hellsberg et al. 2024)), respectively. Residues 86-615 are present in all three protein conformational states. As these simulations were initiated before the publication of substrate-bound structures (Yang and Gouaux 2021), 5-HT^+^ was positioned based on Induced Fit docking with Schrödinger (Hellsberg et al. 2019). Except where noted below, all systems were prepared, equilibrated, and simulated using identical protocols.

### Lipid bilayer optimization

Hydrated protein-POPC systems were optimized using a 50 us-long coarse-grained simulation with Gromacs v2018.8 (Bekker 1993) and the Martini v2.2 force field (Marrink et al. 2007) while constraining the protein secondary structure and internal structure to within ∼1.5Å RMSD for the Cα atoms. The coarse-grained representation efficiently optimizes the protein-lipid interface and reduces equilibration times. A single lipid type was used to minimize artifacts due to the sampling time of rearrangements of more complex mixtures during subsequent atomistic simulations. A representative snapshot with minimal deviation in the positioning of the coarse-grain lipid headgroups (Stix et al. 2023) was converted to all-atom representation (Wassenaar et al. 2014) and the energy-minimized ion-bound protein model (see below) reinserted therein by fitting onto the protein backbone. The resultant periodic box contained ∼150,000 atoms with dimensions of ∼11×11×12 nm. The combined protein-membrane system was energy minimized (5000 steps of conjugate gradients) with positional restraints on protein non-hydrogen atoms and cavity waters with pseudo atoms placed in aromatic rings to prevent lipid trapping, followed by a 500-step minimization with dihedral restraints on backbone and side-chain dihedrals to preserve the protein conformation.

### Ion occupation and protonation

In all systems, the Na1 and chloride sites contained a Na^+^ ion and a Cl^−^ ion, with the latter position based on PDB entry 5I6X. Residue Glu508 was treated as protonated based on proximity to Glu136 and Poisson-Boltzmann calculations of LeuT (Forrest et al. 2007). Residues Cys200 and Cys209 were disulfide bridged, glycosylations were ignored, and truncated termini were capped with neutral groups. Hydrogen atoms in the protein and cavity-bound water molecules added with Dowser (Zhang and Hermans 1996) were energy minimized for 250 steps each using steepest descents and then conjugate gradient protocols. After insertion into the optimized atomistic lipid bilayer described above, complete lipid-protein systems were equilibrated in six MD stages (total >200 ns) while restraining dihedrals of the protein backbone and side chains, and distances for bound Na⁺, Cl⁻, 5-HT^+^ and the Arg104–Glu493 extracellular salt bridge, as detailed below. Protein configurations with different ion combinations or protonation states were each equilibrated separately. Aside from differences in conformational state, we tested the effects of: (a) the Ct2 site populated with Na^+^ or not, and (b) the S1 site in the presence and absence of 5-HT^+^. Asp98 in TM1 interacts directly with the substrate (**Fig. 1B**) and in the absence of substrate, its likely protonation state has not been firmly established. Because empirical or continuum electrostatics calculations rely on static structures and depend on the presence of adjacent ions, we did not attempt to assign a specific pK_A_ and instead explicitly tested both protonation states. Thus, we simulated apo SERT with Asp98 either in its deprotonated state (denoted Asp^−^98) or protonated (denoted AspH98), where the latter corresponds to an elevated pK_A_ of the acidic side chain relative to its intrinsic aqueous value of ∼4.0.

### Parameters for atomistic MD

All-atom simulations were carried out with NAMD v2.14 using the CHARMM36m force field (Huang et al. 2017), parameters of 5-HT^+^ (Huang et al. 2026), and TIP3P waters (Jorgensen et al. 1983). MD simulations were carried out using a 2 fs time step and trajectory output was saved every 4 ps for analyses. A temperature of 298 K was maintained using Langevin Dynamics, and pressure was kept at 1.01325 bar (oscillation time scale = 200 fs; damping scale = 50 fs) using Nosé–Hoover Langevin piston pressure control (Feller et al. 1995). A cut-off distance of 12 Å with a switching function starting at 10 Å was used for Van der Waals interactions. The particle–mesh Ewald method (Darden, York, and Pedersen 1993) was used for long-range electrostatic forces.

### Equilibration restraints

To preclude ion escape or repositioning during equilibration, bound ions were restrained to a minimum number of coordinating interactions, using the COLVARS module (Fiorin, Klein, and Hénin 2013; Fiorin et al. 2024). Specifically, coordinating atoms (Na1 comprising Ala96 O, Asn101 Oδ1, Ser336 O, Ser336 Oγ, Asn368 Oδ1; Ct2 comprising Gly94 O, Val97 O, Leu434 O, and Ser438 Oγ; and the chloride site comprising Asn101 Nδ2, Tyr121 Oη, Gln332 Nε2, Ser336 Oγ, and Ser372 Oγ) were constrained to distances of ≤2.5 Å for Na^+^ ions or ≤3.5 Å for the Cl^−^ ion with a minimum coordination number of 2, thereby preserving the capacity for local rearrangements, and an initial force constant of 10 kcal/mol.

In the outward-occluded conformation, the extracellular pathway salt bridge comprising Arg104 and Glu493 was restrained to ≤3.5 Å (between atoms Arg104 Nε, Nη1 or Nη2 and Glu493 Oδ1 or Oδ2), with a force constant of 50 kcal/mol and a minimum coordination number of 1. When present, 5-HT^+^ was restrained to the S1 site with distances ≤2.8 Å between Asp98 Oδ2 and the 5-HT^+^ cationic nitrogen, as well as Thr439 Oγ1 and the 5-HT^+^ hydroxyl, using a force constant of 200 kcal/mol (Huang et al. 2026).

### Multistage equilibration

During the six equilibration stages, force constants on protein dihedrals and ligand/ion restraints were progressively weakened over 204 ns as described previously (Tan et al. 2022). For the outward-occluded conformation, the Arg104–Glu493 salt bridge restraint was maintained at 50 kcal/mol during the first 4 ns and then reduced stepwise by 10 kcal/mol per stage (Hellsberg et al. 2024). System stability was assessed by monitoring protein backbone RMSD and ion positioning.

### Production MD simulations and analysis

To sample the local impact of each ion and protonation state configuration, production runs were initiated from each equilibrated system but without any restraints applied. Each replica (n=8 for outward-occluded and n=4 for inward- or outward-open states) was initiated with different initial velocities, for ∼0.5 µs, thus precluding major conformational changes. Distances and dihedral angles were calculated with COLVARS module v2022-05-24 (Fiorin et al. 2024) in VMD v1.9.3 (Humphrey, Dalke, and Schulten 1996), using custom wrapper scripts. Calculated data were visualized using python matplotlib v3.9.3 (Hunter 2007). Population distributions were smoothed using kernel density estimation. Secondary structure was analyzed using the MDtraj implementation of DSSP (Kabsch and Sander 1983). This analysis focused on three regions at the cytoplasmic ends of TM4 and TM5 (Fig. 4G): a turn region at the end of TM4 (residues 270-273), the TM4–5 loop (intracellular loop 2; residues 273–277), and the TM5 region adjacent to Thr284 (residues 282–288).

Occupancy maps of water were calculated in VMD using the Volmap plugin v1.1 and visualizations used PyMOL version 3.1.3 (Schrödinger Ltd).

### Whole-cell patch clamping

Whole-cell patch-clamp experiments were performed on HEK293 cells stably expressing SERT or transiently expressing T284A, D437T, and S438A, respectively. HEK293 cells, purchased from ATCC (# CRL-1573; ATCC, USA), were authenticated by STR profiling at the Medical University of Graz (Cell Culture Core Facility). These cells were then used to generate a stable line expressing SERT wildtype or they were transiently transfected with T284A, D437T or S438A, in serum-free DMEM using polyethyleneimine 25kDa (PEI) (Santa Cruz Biotechnology, Inc.; Santa Cruz, TX, USA) as the transfecting agent. The plasmid: PEI ratio was 1:3 (w/w). The cells were regularly tested for mycoplasma contamination by 4′,6-diamidino-2-phenylindole staining. These cells were grown in Dulbecco’s Modified Eagle Media (DMEM) supplemented with 10% heat-inactivated fetal calf serum (FBS), 0.6 µg ml^−1^ penicillin, 1 µg ml^−1^ streptomycin, and 100 μg ml^−1^ of geneticin/G418 for positive selection of transporter-expressing clones. None of the cell lines used in this study were included in the list of commonly misidentified cell lines maintained by the International Cell Line Authentication Committee.

24 hours prior to patching, the cells were seeded at a low density on PDL-coated 35 mm plates. Substrate-induced transporter currents were recorded under voltage clamp. Cells were continuously superfused with a physiological external solution that contained 163 mM NaCl, 2.5 mM CaCl_2_, 2 mM MgCl_2_, 20 mM glucose, and 10 mM 4-(2-hydroxyethyl)-1-piperazineethanesulfonic acid (HEPES) (pH adjusted to 7.4 with NaOH). A pipette solution mimicking the internal ionic composition of a cell contained 163 mM K-mesylate, 1 mM CaCl_2_, 0.7 mM MgCl_2_, 10 mM EGTA (ethylene glycol-bis(β-aminoethyl ether)-N,N,N’,N’-tetraacetic acid), and 10 mM HEPES (pH adjusted to 7.2 using KOH). In experiments in which we omitted intracellular potassium, we used the following solution: 163 mM NMDG-mesylate, 1 mM CaCl_2_, 0.7 mM MgCl_2_, 10 mM EGTA, and 10 mM HEPES (pH adjusted to 7.2 using KOH).

For patch clamp recordings, we employed the whole-cell configuration using an Axon 200B amplifier equipped with an Axon 1550 digitizer (both from Molecular Devices, LLC, San José, CA, USA). Recordings were sampled with 10 kHz. We quantified current amplitudes and associated kinetics using Clampfit software (Molecular Devices, LLC, San José, CA, USA). Passive holding currents were subtracted, and the traces were digitally filtered using an 80 Hz digital 8-pole Bessel low-pass filter.

5-HT or cocaine was applied using a perfusion system (Octaflow II, ALA Scientific Instruments, Inc., Farmingdale, NY, USA) that allowed for rapid and complete solution exchange around the cells within approximately 50 milliseconds. All experiments were conducted at room temperature (25°C) using a temperature control unit (cell microcontrol inline preheater, Green Leaf Scientific, Dublin, Ireland) connected to a temperature control system (PTC-20, npi electronic GmbH, Tamm, Germany). Both the patch-clamp holder and pipette electrode were operated via a PatchStar micromanipulator (Scientifica Ltd., Uckfield, East Sussex, United Kingdom).

### Uptake of [^3^H]5-HT

Radioligand uptake experiments were performed on HEK293 cells transiently expressing wildtype SERT or T284A, D437T, and S438A, respectively. HEK293 cells were seeded on 96-well plates (2 × 10^4^ cells/well) where they were allowed to adhere for 24 hours. Thereafter, the medium was removed, and cells were washed twice with Krebs-HEPES buffer (10 mM HEPES·NaOH, pH 7.4; 120 mM NaCl; 3 mM KCl; 2 mM CaCl_2_; 2 mM MgCl_2_; and 2 mM glucose). Substrate uptake was determined by incubating the cells in a final volume of 0.1 mL of Krebs-HEBES buffer containing 0.2µM [3H]5-HT for 1, 2, 5, 10, 15, 30, and 60 minutes, respectively, at room temperature. Nonspecific uptake was defined in the presence of 10 µM paroxetine. Uptake reactions were terminated by aspiration of the reaction buffer followed by two rapid washes with ice-cold Krebs-HEPES buffer. Subsequently, cells were lysed with 1% SDS to release the retained radioactivity, which was quantified by liquid scintillation counting.

## Results

As a first step in understanding how the presence of a bound cation at the Ct2 site confers conformational change, we characterized the interactions of a Na^+^ ion in this site using MD simulations of holo SERT (i.e., in the presence of 5-HT^+^, Cl^−^, and two Na^+^ ions) in an outward-occluded conformation. We initially focus on an occluded conformation because the interactions at Ct2 are expected to be highly favorable and because using a representation with both pathways closed precludes stochastic ion (un)binding events that complicate the analysis (Hellsberg et al. 2024). As expected from structural and simulation studies of LeuT and other SLC6 transporters (Yamashita et al. 2005; Noskov and Roux 2008; Caplan, Subbotina, and Noskov 2008; Zhao and Noskov 2011; Felts et al. 2014; Zomot, Gur, and Bahar 2015; Ferrandino et al. 2016 ; Tavoulari et al. 2016; LeVine et al. 2018; Szollosi and Stockner 2021, 2022; Hellsberg et al. 2019; Hellsberg et al. 2024; Chan et al. 2022), a Na^+^ ion in the Ct2 site is consistently coordinated by the backbone O atoms of Gly94 and Val97 from TM1 (**Fig. 2A, Fig. S1B**), and frequently also by the backbone O atom of Leu434 from TM8 (**Fig. 2A-B, Fig. S1B**). At least two water molecules are within coordination distance of the Na^+^ ion (**Fig. S2C**). Lastly, two side chains from TM8 are consistently involved: a carboxylate oxygen from Asp437 and the hydroxyl oxygen of Ser438 (**Fig. 2A-B**), in line with observations that these residues are critical for major conformational changes in SERT (Tavoulari et al. 2016).

**Figure 2.**
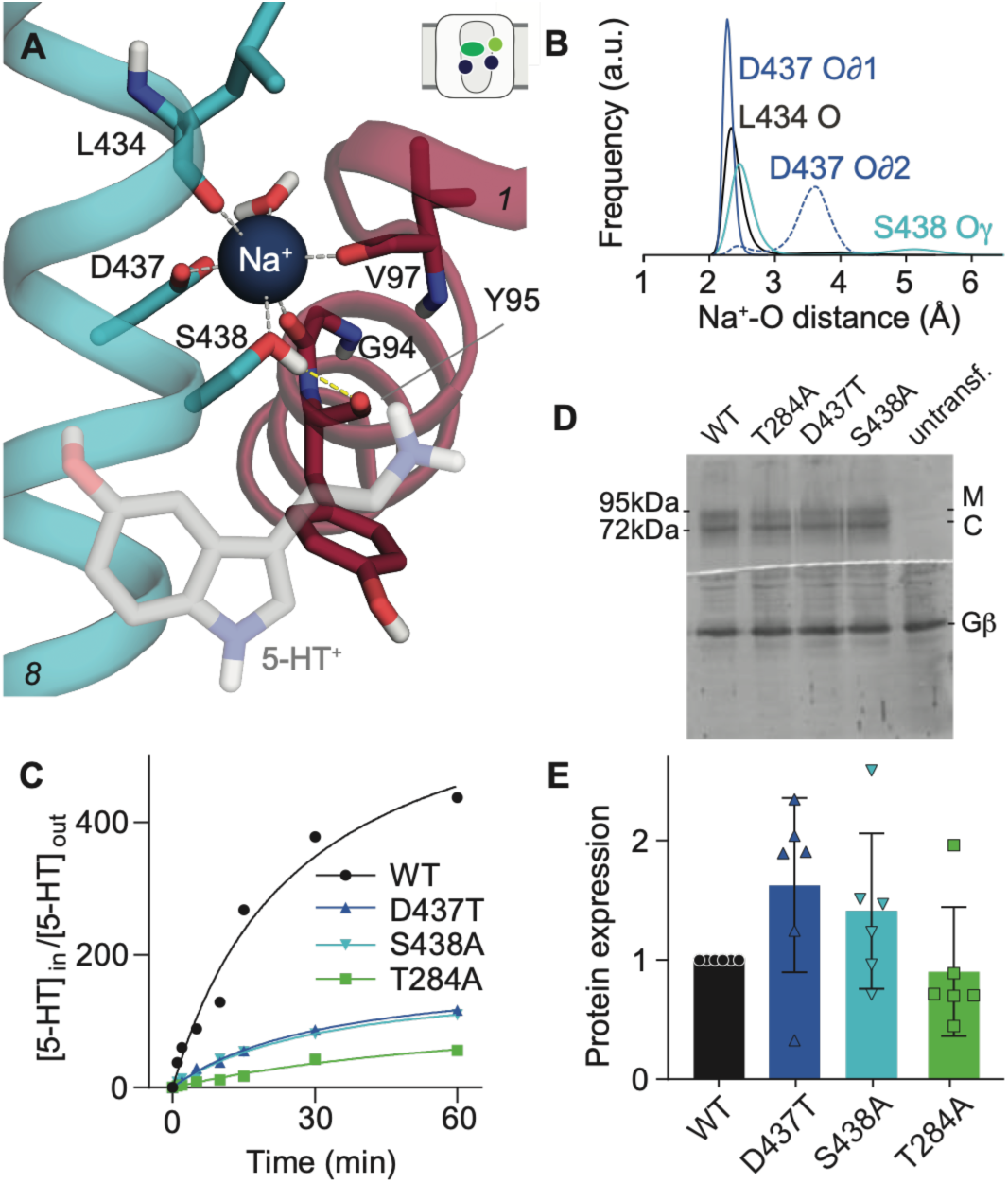
Interaction network including the Ct2 site of SERT. **(A)** Molecular dynamics simulation snapshot of the outward-occluded state of SERT bound to 5-HT^+^, showing TM1 (*red*) and TM8 (*blue*), with Ser438 in the *gauche*(+) orientation. 5-HT^+^ is shown as transparent sticks. Side chains and water are shown as sticks, and Na^+^ is shown as a blue sphere. A second water molecule is obscured by the Na^+^ ion. Only polar H atoms are shown explicitly, for clarity. Inset: schematic indicating the occluded conformation with bound two sodium (*blue circles*), chloride (*light green circle*) and 5-HT^+^ (*green oval*). **(B)** Na^+^ coordination distances of O atoms in TM8 during MD simulations of the outward-occluded state. Distances are shown as a distribution over data collected from n = 8 trajectories, each 0.5 µs long. Distances were measured to the backbone O of Leu434 (*black line*), the two O∂ oxygen atoms of Asp437 (*blue solid and dashed lines*, respectively), and the hydroxyl O atom of Ser438 (*cyan line*). **(C)** Uptake of [^3^H]-5-HT^+^ into cells expressing SERT wild type (*black circles*), D437T (*blue up triangles*), S438A (*cyan down triangles*), and T284A (*green squares*). The concentrative power ([5-HT]_in_/[5-HT]_out_) is plotted as a function of the exposure time. Cells were incubated with 200 nM of [^3^H]-5-HT^+^ for 1, 2, 5, 10, 15, 30, and 60 minutes. Each experiment was conducted in triplicate, with each data point representing the mean of three independent experiments. The lines are fits of a mono-exponential function to the data points. The rates estimated by the fit were: wild-type (0.048 ± 0.008 min^-1^), D437T (0.037 ± 0.011 min^-1^), S438A (0.038 ± 0.006 min^-1^), and T284A (0.020 ± 0.012 min^-1^). (**D**) Western blot showing relative expression levels of SERT wild-type and Ct2 mutants. Membrane proteins extracted from these cells were denatured and resolved with SDS-PAGE and transferred onto nitrocellulose membranes. Blots were incubated overnight at 4°C with anti-GFP (*top*) or anti-Gβ (*bottom*, loading control) antibodies. The immunoreactive bands were detected with fluorescently labeled secondary antibodies. The blot is representative of six experiments. The core-glycosylated band (C) and the mature glycosylated band (M) levels are indicated. **(E)** Relative protein expression levels (i.e., density value of M / density value of Gβ normalized to the same ratio obtained for wild-type, *black*) for D437T (1.65 ± 0.62, *blue*), S438A (1.41 ± 0.55, *cyan*), and T284A (0.90 ± 0.46, *green*) from n = 6 Western blots.

### Ct2 site residues are important for coupling efficiency

Patch clamp traces recorded for SERT mutants modified at the Ct2 site, namely D437T and S438A, unexpectedly exhibited a current that was present even without applying substrate (Hellsberg et al. 2024). Although the origins of this constitutive current remain to be established, its presence suggests a substrate-independent flux (“leak”) of ions and raises the possibility that the mutated transporters have reduced coupling efficiency. To determine whether coupling efficiency is affected in these mutants, we estimated the concentrative power by measuring substrate uptake over time. Specifically, we compared the uptake of 0.2 µM [^3^H]-5-HT^+^ by wild-type SERT, D437T, and S438A. Cells expressing the Ct2-site mutants accumulated less 5-HT^+^ than cells expressing wild-type SERT (**Fig. 2C**). Additionally, we found that cells transiently expressing YFP-tagged wild-type SERT, D437T, and S438A expressed similar levels of both the ER-resident core-glycosylated protein and the transporter harboring mature glycans (bands labeled C and M, respectively in **Fig. 2D**). Thus, if anything, when corrected for the levels of a reference marker, the G protein β-subunit (Gβ; **Fig. 2D**), the quantitative data suggested that there were higher levels of D437T and S438A than of wild type SERT (**Fig. 2E**). This finding, together with our understanding that steady state transport levels are largely insensitive to cell surface expression levels, suggests that the reduced steady state 5-HT^+^ uptake reflects impaired concentrative power in the mutants, consistent with the Ct2 site being critical for Na^+^-substrate coupling.

### Analysis of Asp437 reveals interactions with TM5

Structural data suggest Na^+^ binding at Ct2 promotes the apposition of TM1 and TM8, a hallmark of cytoplasmic pathway closure. However, this TM1-TM8 interface is not exclusive to the Na^+^-bound state; it is also a structural requirement for the transitions that reset the transporter to the outward-open state. Thus, the action of Na^+^ coordination to bring together TM1 and TM8, while essential for serotonin uptake, cannot be solely responsible for all functionally important conformational responses of the transporter. We posit that the second coordination shell of the Ct2 site plays a role in transducing binding to shifts in the global conformational ensemble; accordingly we carried out detailed analysis of the interaction network during the MD simulations of the outward-occluded state. The Na^+^-Asp437 interaction remained constant on the sub-microsecond timescale (**Fig. 2B; Fig. S1B**), with one O∂ atom in the carboxyl group coordinating the Na^+^ ion (**Fig. S1B**). Notably, the other O∂ atom accepts a hydrogen bond from a water molecule or from the hydroxyl of Thr284 (**Fig. 3A**). Thr284 is in TM5, whose cytoplasmic end unwinds significantly when adopting the inward-open state of SERT (Coleman et al. 2019; Yang and Gouaux 2021). We therefore investigated any interactions involving Thr284 that might underlie the transporter’s conformational dependence on Na^+^ binding.

**Figure 3.**
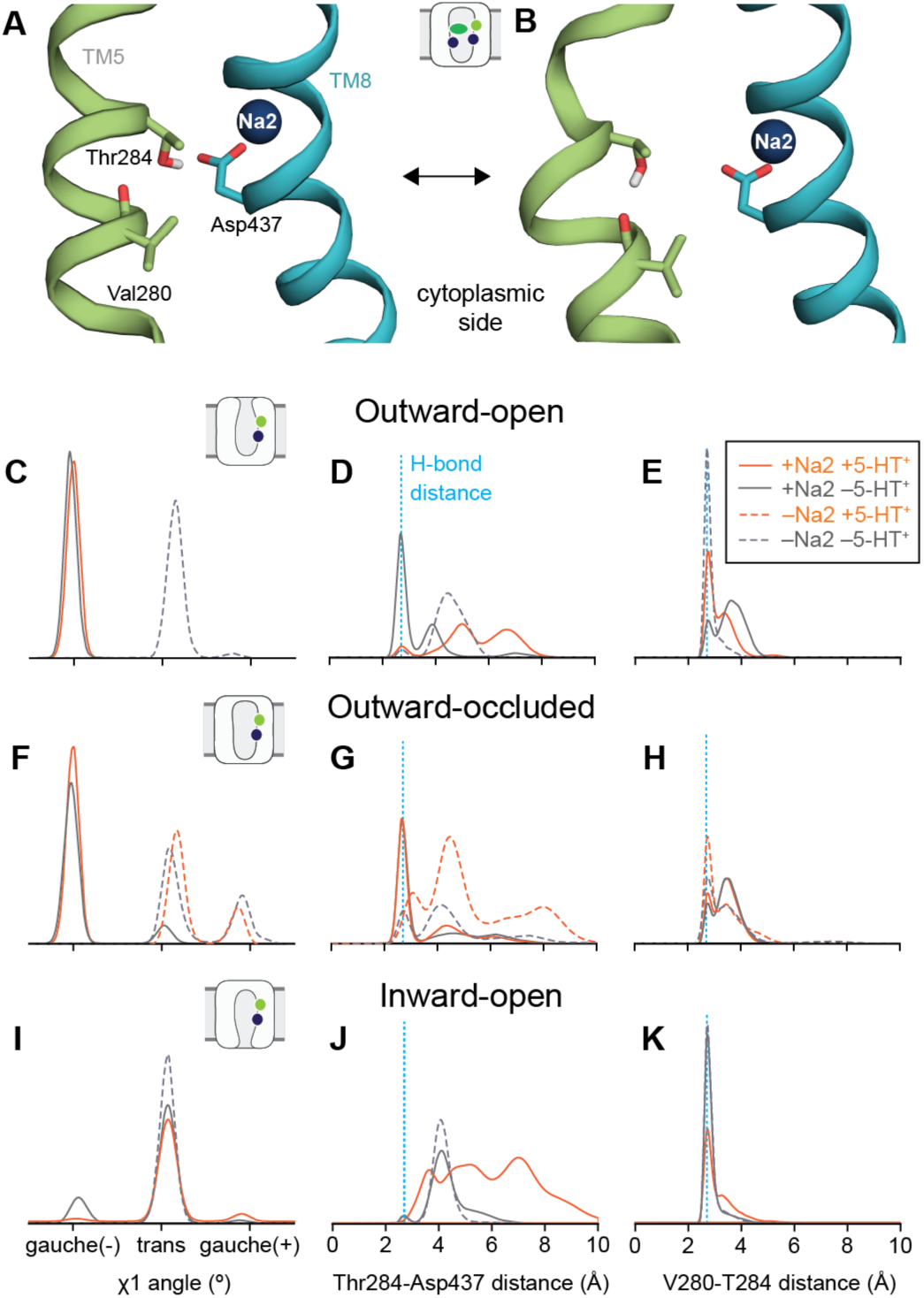
State- and ligand-dependent interactions within the Ct2–TM5 network of SERT. **(A–B)** Simulation snapshots of the occluded state of SERT illustrating the two competing hydrogen-bond configurations of Thr284 observed in simulations: donation to the carboxyl group of Asp437 (A) or to the backbone carbonyl of Val280 (B); the Na^+^ ion at Ct2 is shown as a blue sphere. **(C-K)** Distributions of the Asp437 side-chain χ1 dihedral angle (C,F,I), the distance between the Thr284 hydroxyl O atom and the closest carboxyl O atom of Asp437 (D,G,J), and the distance between the Val280 backbone carbonyl O atom and the Thr284 hydroxyl O atom (E,H,K), from simulations of outward-open (C-E), outward-occluded (F-H) and inward-open (I-K) SERT, as indicated by the schematics on the right. Simulations were performed with (*solid lines*) or without (*dashed lines*) Na^+^ at the Na2 site. The S1 site was either empty with Asp98 deprotonated (*gray*) or initially occupied with 5-HT^+^ (*orange*). For the inward-open conformation, a Na^+^ ion initially placed at Ct2 readily moved away. The distance corresponding to a typical O–O hydrogen bond (2.7 Å) is indicated by a vertical dashed line. All distributions are computed over n = 8 or n = 4 independent 500-ns trajectories. Corresponding time-resolved data are shown in **Fig. S3**.

### Thr284 interactions with Na^+^ bound

An adaptive sampling simulation study of SERT indicated that dissociation of the Ct2 site Na^+^ ion is coupled to reorientation of the Asp437 side-chain (Chan et al. 2022). Accordingly, we asked whether Na^+^ binding at the Ct2 site stabilizes specific Asp437 side-chain orientations that in turn influence the Thr284 interaction network, and we analyzed the behavior of these residues in additional MD simulations of SERT. Before examining how these interactions vary across conformational states, we first further characterized the outward-occluded state, where Na^+^ coordination at Ct2 is likely the most complete and stable. In the occluded state with both Na^+^ at Ct2 and 5-HT^+^ in the S1 site, the Asp437 side chain predominantly adopted a *gauche*(–) χ_1_ angle (**Fig. 3F**, *orange*). This orientation is compatible with the interhelical hydrogen bond from Thr284 to Asp437 (**Fig. 3A; 3G***, orange*). However, Thr284 also formed an intrahelical H-bond by donating its side chain hydroxyl hydrogen to the backbone carbonyl of Val280, located one helical turn away on the cytoplasmic side of TM5 (**Fig. 3B**), resulting in a peak in the Val280-Thr284 distance distribution at around 2.7Å (**Fig. 3H**, *orange*). These arrangements were largely independent of the presence or absence of 5-HT^+^ in the occluded S1 site (**Fig. 3F-H***, solid lines*).

To assess the interdependence of the two interactions of Thr284 (i.e. with Val280 or Asp437) we plotted the corresponding distances over simulation time (**Fig. S3B**). Periods in which the Thr284-Asp437 distance was increased coincided with H-bonding between Thr284 and Val280 and direct comparison of the two distances gave a mean Spearman coefficient of –0.40 ± 0.06, consistent with anti-correlated, thermally-fluctuating interactions. Thus, the Thr284 hydroxyl alternated interactions between TM8 and its neighboring backbone in TM5.

Because the simulations presented so far reflect a single (occluded) conformation of SERT, it remains unclear whether the Thr284-Asp437 interaction varies with transporter conformation. We therefore carried out simulations initiated with Na^+^ at the Ct2 site for both outward-open (Huang et al. 2026) and inward-open SERT (Hellsberg et al. 2019) and analyzed the Thr284 network therein. In the inward-open state, cations placed near the Ct2 site rapidly unbound and entered the aqueous solution during the equilibration stages whether or not 5-HT^+^ was bound (**Fig. S1C**). This lack of consistent Na^+^ coordination was associated with Asp437 rarely visiting the *gauche*(–) orientation (**Fig. 3I**; **Fig. S3C, S3F**). Furthermore, Thr284 and Asp437 were too distant to form an H-bond (**Fig. 3J**; **Fig. S3C, S3F**), and Thr284 instead tended to interact with its backbone (**Fig. 3K**; **Fig. S3C, S3F**). In other words, the TM5-TM8 interaction is broken during simulations of the inward-open state, independent of the presence of 5-HT^+^ at the S1 site, or a Na^+^ ion placed initially next to Asp437.

In the outward-open state with a Na^+^ ion at Ct2, Asp437 adopted the same *gauche*(–) orientation (**Fig. 3C***, solid lines*) seen in the occluded state. In addition, the H-bonding interaction between Asp437 and Thr284 was observed in the outward-open state, but only in the absence of 5-HT^+^ (**Fig. 3D**, *gray*). Thus when the extracellular pathway is open, binding of 5-HT^+^ disfavors the Asp437-Thr284 H-bond (**Fig. 3D**, *orange*). A clue to the origins of these different behaviors could be found in the reduced Na^+^ coordination by Ser438 (**Fig. S1A, Fig. S4A**), due to the primary amine of 5-HT^+^ providing an alternate interaction partner when Ser438 adopts the *gauche*(–) orientation (**Fig. S4D**). To compensate for the lack of Ser438 interactions, the Ct2 Na^+^ ion tended to engage both oxygen atoms of Asp437 (**Fig. S1A**) and this divalent ion coordination was associated with longer distances to Thr284 (**Fig. 3D-E***, orange*; **S3A***, reds*). Thus, with the extracellular pathway open, the positions of 5-HT^+^ and Na^+^ were less compatible with stabilizing the TM5-TM8 H-bond. The suboptimal Thr284 interaction network seen for the holo outward-open state is however likely to be short-lived, because binding of 5-HT^+^ enables occlusion of the extracellular pathway on the microsecond timescale (Gradisch et al. 2022), after which the Thr284 network would be re-established. Nevertheless, the simulation data for both inward- and outward-open states support the notion that optimal coordination of the Ct2 Na^+^ ion favors the Asp437 and Thr284 H-bond and corresponding close proximity between TM5 and TM8.

### Analysis of Na^+^-free states

To directly assess the contribution of the Na^+^ ion, we performed simulations of the same states but without a Na^+^ ion initially at Ct2. Notably, in both outward-open and outward-occluded states, i.e. with the cytoplasmic pathway closed, the charged Asp437 in the Ct2 site appeared to attract a small number of water molecules along the interface between TM5 and TM1/TM8 from the cytoplasm up to the Na^+^ ion (**Fig. 4A-D**). In one simulation without Na^+^ bound, this hydrated pathway allowed repeated Na^+^ ion permeation from the salt solution to Asp437 (**Fig. 4E**). Nevertheless, in the Na^+^-free condition, the Asp437 rotamer and Thr284 interaction distributions were shifted significantly (**Fig. 3A-H***, dashed lines*; **Fig. S3D-E**), away from the Na^+^-bound peaks (**Fig. 3A-H**, *solid lines*) and toward those seen in the unbound, inward-open state (**Fig. 3I-K; Fig. S3C, S3F**). Specifically, in the absence of a Na^+^ ion, Asp437 adopted a trans orientation and rarely H-bonded with Thr284. Thus, the simulation data suggest that Na^+^ at the Ct2 site stabilizes the specific Asp437 arrangement that favors interhelical hydrogen-bonding with Thr284 raising the possibility that Thr284 plays a key role in modulating the gating of TM5 in SERT.

**Figure 4.**
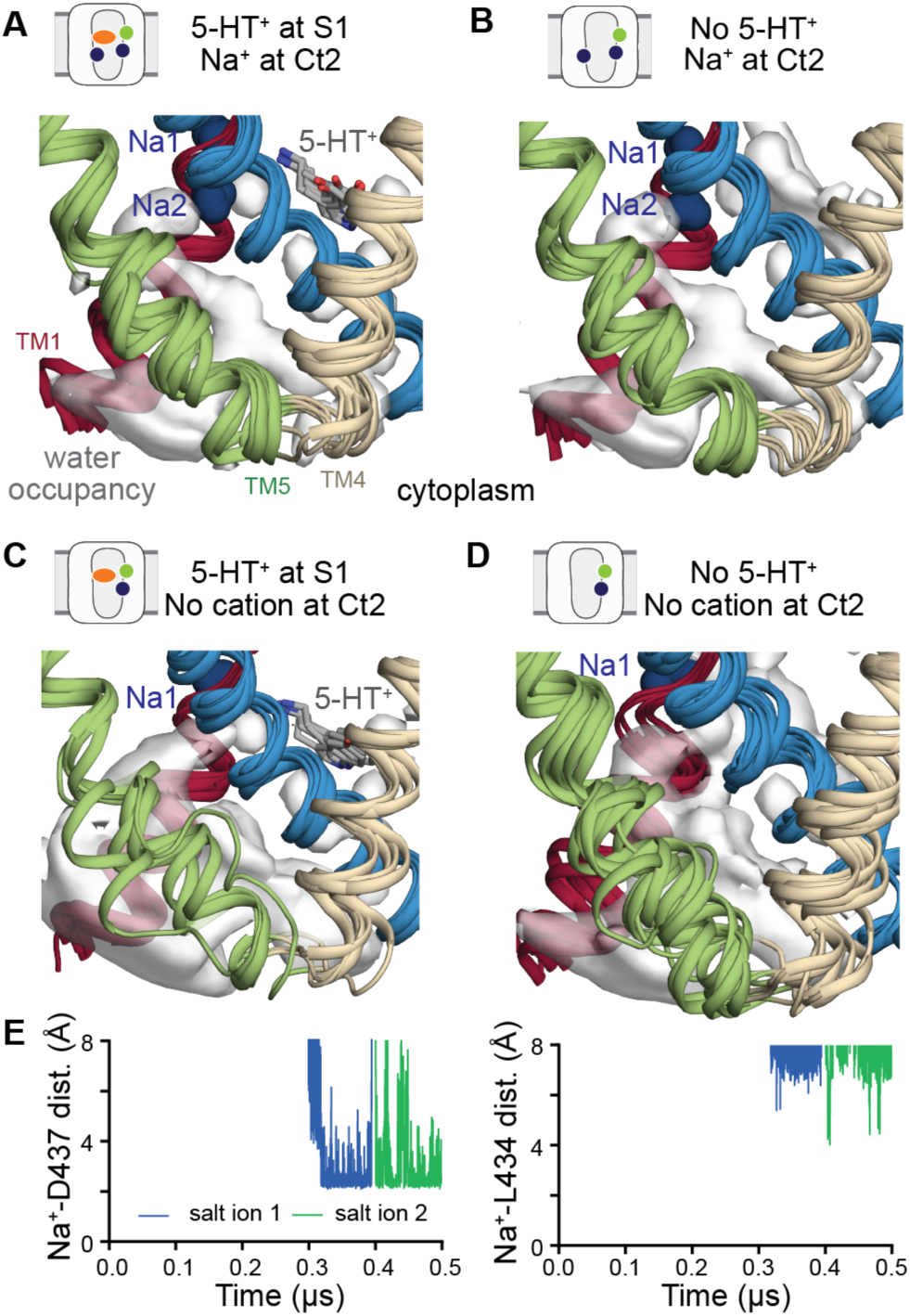
Dynamic behavior of the cytoplasmic end of human SERT in the outward-occluded conformation. **(A-D)** Structural snapshots of the TM4-5 loop region superposed on water occupancy maps (*white volume*) from MD simulations of outward-occluded SERT, with the Ct2 site occupied by Na^+^ (*blue sphere,* A-B) or without a cation (C-D) as indicated. The S1 site contained 5-HT^+^ (*gray sticks;* A, C), or no ligand (B, D). Helices TM1a (*red*) and TM8 (*blue*) are shown for reference alongside TM4 (*wheat*) and TM5 (*green*). The uppermost blue sphere in each panel is the Na1-site sodium ion. Each panel shows the final snapshot from each of four or eight independent 500 ns-long trajectories. **(D)** Spontaneous Na^+^ ion penetration from the cytoplasmic side toward Asp437 during simulations of outward-occluded SERT. The Ct2 site was initially unoccupied by cations, and the S1 site contained 5-HT^+^. The distance between individual Na^+^ ions from the cytoplasmic salt solution and either the carboxylate oxygen of Asp437 (*top*) or the backbone oxygen of Leu434 (*bottom*) is shown as a function of time.

### Thr284 is important for Na^+^ coupling efficiency

To assess the predicted role of Thr284, we tested whether this residue contributes to sodium-dependent substrate coupling in cells. T284A exhibited a markedly reduced concentrative power relative to both WT and the Ct2 mutants D437T and S438A (**Fig. 2C**). This reduction supports a role for Thr284 in efficient Na^+^-coupled transport, consistent with the predicted requirement for a side chain hydroxyl at this site.

To investigate how the hydroxyl group of Thr284 helps couple Na^+^ transport to 5-HT^+^ transport, we examined whether its interaction with Asp437 is associated with changes in TM5. We focused on two features: local unwinding near Thr284 and loss of helical structure in the cytoplasmic TM4-5 loop (**Fig. 5A**).

**Figure 5.**
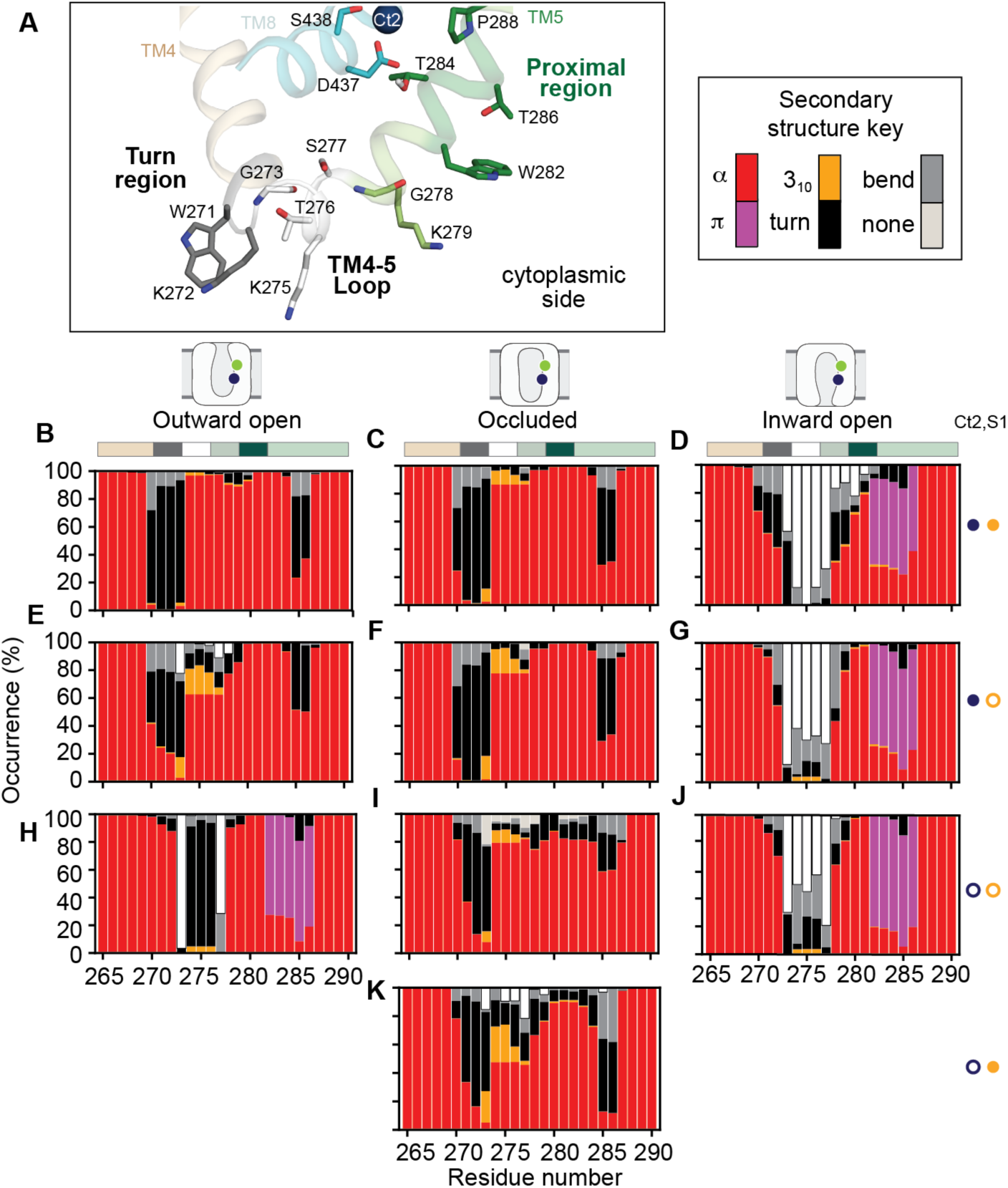
State- and ligand-dependent secondary structure of the TM4-5 region of human SERT. **(A)** Example structure of the TM4–TM5 region, illustrating the regions analyzed. The turn region at the end of TM4 (residues 270–273, *gray*) precedes the TM4-5 loop (residues 273-277, *white*). The TM5 region proximal to the Ct2 coupling residue Thr284 (residues 282–286; *dark green*), lies adjacent to the Ct2 Na^+^ binding site. Residues involved in Ct2 Na⁺ coordination on TM8 (Asp437 and Ser438; *cyan*) and the Ct2 ion (*blue sphere*) are shown for reference. Key residues are displayed as sticks, and backbone atoms are omitted where possible for clarity. **(B-G)** Stacked bar plots show the mean percentage occupancy of DSSP-assigned secondary structure states (α-helix, 3_10_-helix, turn, bend or no secondary structure) for the TM4-TM5 region (replica-averaged, *n* = 4 (panels B, D, G, J, K), n = 8 (panels C, F, I), and *n* = 1 (panel H)). The protein is in an outward-open (left), outward-occluded (middle) or inward-open (right) conformation, as indicated by the schematics above B-D. Simulations were performed with the Ct2 site occupied by Na^+^ (filled blue circle, top rows) or no cation (open blue circle, bottom rows) and with the S1 site occupied by 5-HT^+^ (filled orange circle, top and bottom rows) or empty (open orange circle, middle rows). Analyses focus on residues 265-290, which includes the turn region in TM4, the TM4-5 loop, and the TM5 region near Thr284, as defined in panel G and indicated by horizontal, colored bars. Occupancies summarize time-dependent DSSP assignments across the full trajectory length; corresponding time-resolved DSSP data for individual trajectories are shown in **Fig. S4.**

### MD simulations characterize the secondary structure of TM5

In simulations with Na^+^ bound initiated from the outward-open and outward-occluded structures, the TM4-5 loop remained predominantly α-helical throughout the sampled trajectories (**Fig. 5B-C, 5E-F**; **Fig. S5A-B, S5D-E**). In simulations initiated from the inward-open structure, this loop consistently lacked a defined secondary structure (**Fig. 5D, 5G**; **Fig. S5C, S5F**).

Notable differences were also seen in TM5 near Thr284. In the inward-open state, residues 282-286 primarily sampled an π-helical arrangement, a more loosely wound arrangement than an α-helix. (**Fig. 5D, 5G**). This region rapidly switched between π-helical, α-helical and non-helical turn or bend conformations (**Fig. S5C, S5F**). In the outward-open and outward-occluded Na^+^- bound simulations, the corresponding region did not form this π bulge. Instead it adopted α-helical α-helical or turn conformations (**Fig. 5B-C, 5E-F**; **Fig. S5A-B, S5D-E**).

For the outward-occluded and inward-open states, TM5 showed similar features both with and without 5-HT^+^ present (**Fig. 5C-D** vs **5F-G**). Thus, within these simulations, 5-HT^+^ had little effect on TM5 secondary structure once the extracellular pathway was closed. In the outward-open state, however, residues 285 and 286 were slightly more likely to be α-helical when 5-HT^+^ was absent. This observation is consistent with a model in which greater helicity helps stabilize the closed cytoplasmic pathway before 5-HT^+^ binding, thereby limiting Na^+^ leakage.

*Na^+^ removal favors local unwinding near Thr284 on TM5* – We next compared simulations with and without Na⁺ at Ct2 to ask how site occupancy affects the secondary structure sampled from each starting conformation. In the occluded state, removing Na^+^ did not result in a complete conversion to a π-helix. Nevertheless, several simulation replicas showed reduced α-helicity in the residues preceding the segment near Thr284 (**Fig. 5I, 5K**). These residues lost the backbone H-bonds characteristic of an α -helix, which connect residues four positions apart (*i* to *i*+4) and adopated a turn conformation (**Fig. S5H, S5J**). This change is consistent with an initial step toward π-helix formation.

In the Na^+^-free outward-open simulation, this segment of TM5 near Thr284 rapidly converted to the π-helix. However, only one replica was obtained for this condition: the open extracellular pathway readily allowed Na^+^ ions to enter the binding site (**Fig. S4B**), making a persistently Na^+^- free state difficult to maintain. This result should therefore be interpreted cautiously.

Together, these observations suggest that Na^+^ binding stabilizes Asp437 and its H-bonding with Thr284, restricting local unwinding of TM5. When Na^+^ is absent, the Thr284-Asp437 interactin is lost, and TM5 can separate from TM8. We propose that this separation permits further unwinding near Thr284 and formation of the backbone H-bonding characteristic of a π-bulge, which connects residues five positions apart (*i* to *i*+5).

The cytoplasmic TM4-5 loop showed a different response. In the outward-occluded and inward-open states, removing Na^+^ did not markedly change its secondary structure relative to the corresponding Na^+^-bound simulations (**Fig. 5I-K**; **Fig. S5H-J**). Thus, Na^+^ removal alone did not substantially alter the loop’s structure in these states. We propose that full unwinding of this loop requires sufficient separation of TM5 from TM8.

### The Thr284–Asp437 interaction may link Na⁺ occupancy to TM5 packing

Our analysis characterize individual conformational states at equilibrium; they do not directly establish the sequence of transitions between them, which would require enhanced sampling strategies. Nevertheless, results support a model of mechanistic coupling in which Thr284 links Na^+^ occupancy at Ct2 to the structure and packing of TM5.

In this model, Na^+^ unbinding from the fully occluded state through the small hydrated channel leading into the cytoplasm allows Asp437 to rotate freely. This reduces its propensity to interact with Thr284, allowing TM5 to move away from TM8. The resulting loss of structural restraint permits local unwinding near Thr284 and, subsequently, unwinding of the cytoplasmic loop. This model predicts that mutation of either Asp437 or Thr284 would impair the ability of TM5 to respond to Na^+^ occupancy and maintain the close packing needed to prevent Na^+^ release before 5-HT^+^ captured.

### Probing the role of Ser438 in coupling

The mutational analysis shows that Ser438 also contributes to the concentrative power of SERT. Its position on the extracellular side of the Na^+^ ion at Ct2 suggests a role distinct from that of Asp437. To investigate this role, we examined the interactions sampled by Ser438 during the MD simulations.

### Ser438 samples two alternative interaction patterns

In simulations initated from the outward-occluded structure with Na^+^ bound at Ct2, Ser438 predominantly adopted an orientation that connected the ion-binding site to TM1. In this orientation, its hydroxyl oxygen directly coordinated the Na^+^ ion (**Fig. 2A; Fig. S1B**), and its side-chain χ_1_ torsion angle was primarily in the *gauche*(+) conformation (**Fig. 6B**). The hydroxyl group also donated a hydrogen bond to the backbone oxygen of Tyr95 in TM1a (**Fig. 2A**, **Fig. 6C**)(Chan et al. 2022). Thus, during these simulations, Ser438 directly linked TM8 to TM1. This interaction could help stabilize closure of the cytoplasmic pathway. In this orientation, the Ser438 side chain also occupied the space between the Ct2 and S1 pockets (**Fig. 2A**), consistent with its proposed role in controlling movement between these sites (Felts et al. 2014; Szollosi and Stockner 2021). This arrangement may therefore also restrict Na^+^ movement from Ct2 toward S1, and ultimately, back to the extracellular solution.

**Figure 6.**
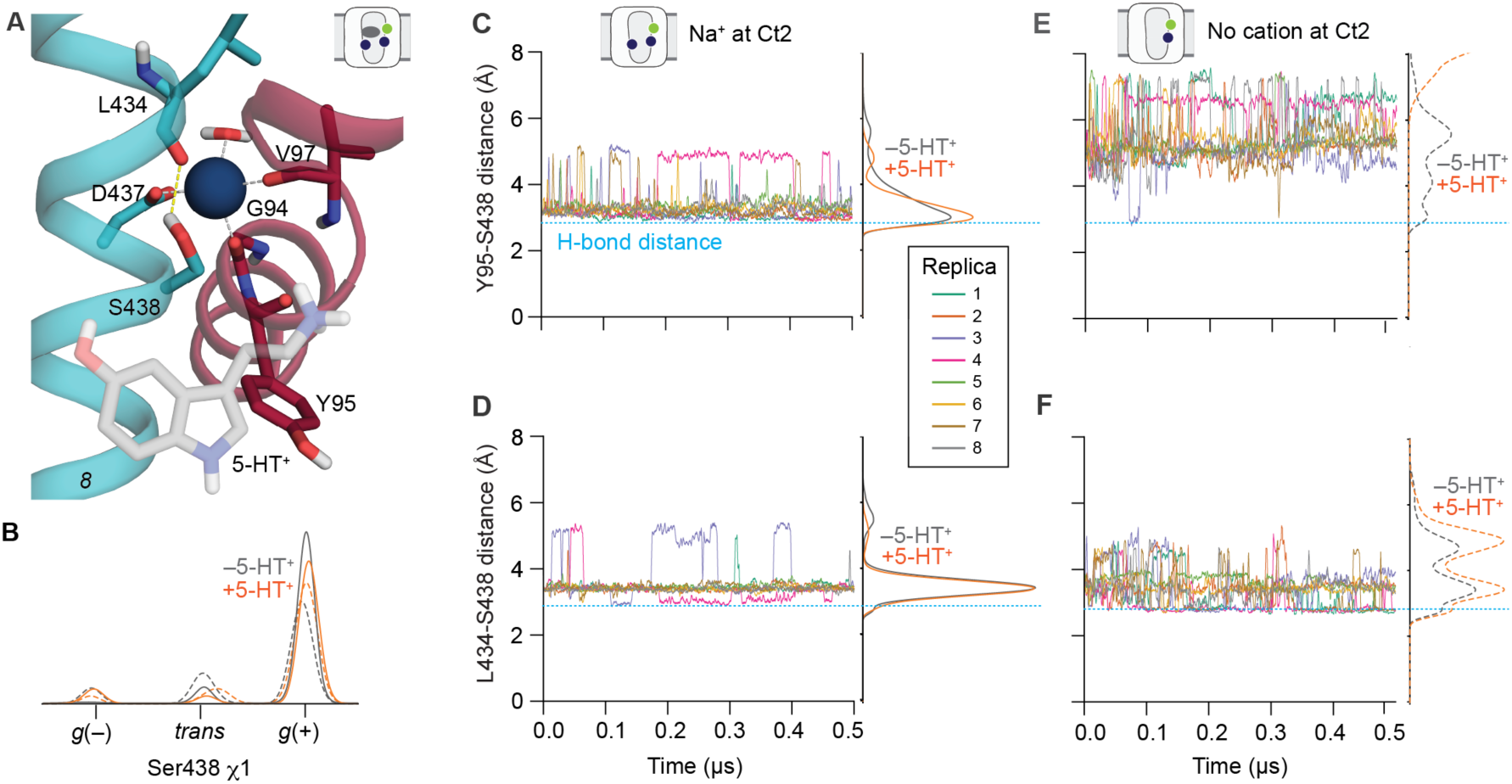
Ser438 interactions with the backbone of TM1 in the outward-occluded state. **(A)** MD simulation snapshot of the outward-occluded state of SERT bound to 5-HT^+^ showing TM1 (*red*) and TM8 (*cyan*) with Ser438 in the *gauche*(–) orientation. See Fig. 2A for analysis of the *gauche*(+) orientation and additional descriptions. **(B)** Distribution of the side chain χ_1_ dihedral angle of Ser438 over *n* = 4 or 8 trajectories of the outward-occluded state, in the presence (*solid*) and absence (*dashed*) of a Ct2-bound Na^+^ ion. Simulations were carried out with the S1 site occupied by 5-HT^+^ (*orange*), or no ligand (Asp98 deprotonated; *gray*). **(C-F)** Distances from the side chain oxygen of Ser438 to the backbone carbonyl oxygen of either Tyr95 in TM1a (C,E) or Leu434 in TM8 (D,F) as a function of trajectory time for each replica in the 5-HT^+^-free system, colored according to the replica legend. Distributions of the data for both 5-HT^+^ bound (*orange*) and free (*gray*) states are shown to the right of each plot. The Ct2 site contained Na^+^ (C-D, *solid distribution lines*) or no cation (E-F, *dashed distribution lines*) as indicated in the schematics. In each case, *n* = 4 or 8 trajectories were computed. The distance of a typical O-O H-bond (∼2.7 Å) is indicated with a horizontal dashed line.

Ser438 occasionally switched to a second interaction pattern. In a small population of trajectory frames, the hydroxyl oxygen moved >5 Å from Na^+^ (**Fig. 2B; Fig. S1B**) and the side-chain adopted the alternative *gauche*(–) orientation (**Fig. 6A-B**). In this configuration, Ser438 formed a hydrogen bond within TM8, with Leu434 one helical turn away, and lost its interaction with Tyr95 in TM1a (**Fig. 6D**). In equivalent simulations of the outward-open state with both Na^+^ and 5-HT^+^ bound, the Ser438-Tyr95 interaction was also favored (**Fig. S4E**), but with a shift toward a higher population of alternate conformations (**Fig. S4D**). The simulations therefore suggest that the Na^+^-associated connection between TM1 and TM8 mediated by Ser438 is stabilized in the outward-occluded conformation, in which the extracellular pathway is closed.

### Na⁺ occupancy favors the Ser438–Tyr95 interaction

To test the dependence of these interactions on Na^+^ occupancy, we compared simulations of the outward-occluded state with and without Na^+^ at Ct2. Without Na^+^, the Ser438-Tyr95 interaction was no longer observed (**Fig. 6E**) and the Ser438-Leu434 hydrogen bond became more prevalent (**Fig. 6F**). A similar shift was observed for simulations of the outward-open state without Na^+^ (**Fig. S4E-F**; *dashed line*), and the resulting arrangement reflected that of the inward-open state, where the Ser438 side chain was more than 5Å from the backbone of Tyr95 (**Fig. S4H-I**). By contrast, removing 5-HT^+^ from S1 while retaining Na^+^ at Ct2 did not appreciably change the Ser438 interaction pattern relative to the holo transporter (**Fig. 6C-D**). In these simulations therefore the interaction pattern depended on Na^+^ occupancy at Ct2 rather than on 5-HT^+^ occupancy at S1.

Loss of the Ser438-Tyr95 interaction was accompanied by increased separation of the residues themselves. With Na^+^ bound at Ct2, the mean distance between their backbone Cα atoms was ∼4.2 Å regardless of 5-HT^+^ occupancy. Without Na^+^, this distance exceeded 6Å. Thus, the change in interaction pattern involved both side-chain reorientation and a local change in backbone separation at Ser438 and Tyr95.

### Two interaction networks may link Ct2 occupancy to coupling

Together, the simulations support a model in which Ct2-bound Na^+^ influences two interaction networks (**Fig. S6**). Through Asp437 and Thr284, Na^+^ occupancy is linked to the local winding and position of TM5. Through Ser438 and Tyr95, it is linked to the arrangement of the region between Ct2 and S1. We proposed that these complementary networks help coordinate structural rearrangements on the cytoplasmic and extracellular sides of Ct2, supporting efficient coupling of Na^+^ and 5-HT^+^.

### Electrophysiology measurements support a role for Thr284 in Na^+^ coupling

Our simulations predict that efficient coupling between Na^+^ and 5-HT^+^ uptake requires not only the two Ct2 site residues Asp437 and Ser438, but also Thr284 in TM5, which links Ct2 Na^+^ occupancy to cytoplasmic conformational changes. Consistent with this prediction, whole cell patch clamp recordings of the Ct2 mutants D437T and S438A (Hellsberg et al. 2024) revealed that they both mediate a pronounced, substrate-independent, constitutive current through SERT supporting a role for these residues in coupling beyond merely recognizing Na^+^. To better define the roles of these residues in the energetics of coupling, we used patch clamp recordings to extend the electrophysiological analysis of D437T and S438A as well as to characterize the T284A mutant.

Application of a saturating concentration of 5-HT^+^ (30 µM) elicited an inwardly directed peak current in wild-type SERT (**Fig. 7A**), and the Ct2-site mutants D437T (**Fig. 7B**) and S438A (**Fig. 7C**), but not in the TM5 coupling mutant T284A (**Fig. 7D**). The inward peak current reflects a transport-associated conformational transition that allows 5-HT^+^ and Na^+^ to be released into the cytoplasm (**Fig. 1A**). T284A lacks the transport-associated peak current, suggesting that removal of the Thr284 side chain impairs engagement of the TM5-dependent conformational transition required for opening the cytoplasmic pathway.

**Figure 7.**
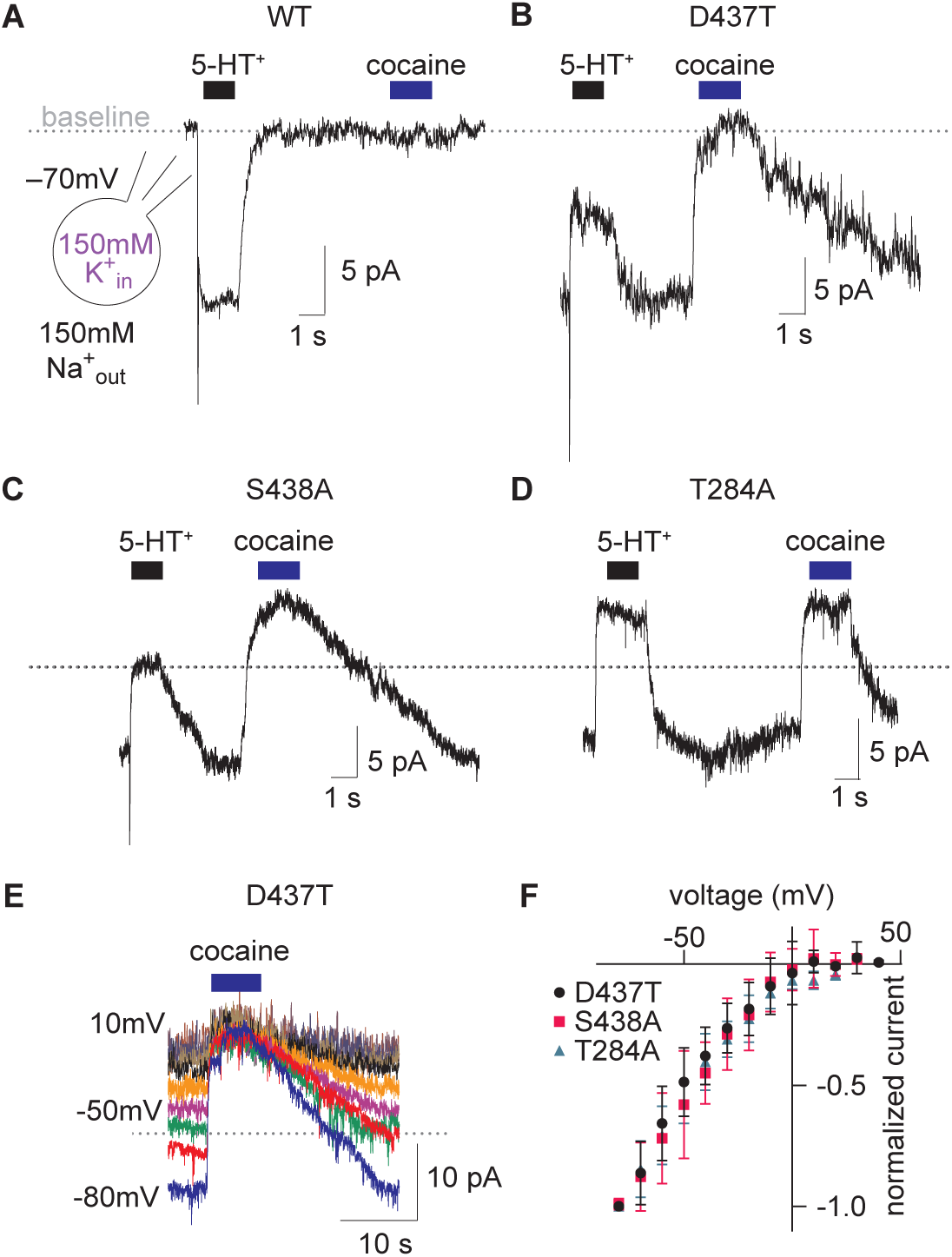
Ct2 pathway mutants mediate a constitutive uncoupled current. (**A**) 5-HT^+^ (30 µM) induced current obtained from a cell expressing wild-type SERT recorded at the conditions illustrated in the inset. The blue bar indicates the subsequent application of a saturating concentration of cocaine (i.e., 30 µM). (**B**) Current obtained from a cell expressing D437T recorded at the same conditions as in (A). The application of 30 µM 5-HT^+^ induced a peak current and a subsequent upward deflection of the current. Administration of 30 µM cocaine also gave rise to an upward deflection of the current but the amplitude thereof was greater than that induced by 5-HT^+^. (**C**) Current recorded from a cell expressing S438A. The current mediated by this mutant phenocopied that of D437T. (**D**) Current obtained from a cell expressing T284A. This mutant lacked the 5-HT^+^-induced peak current but also showed an upward deflection of the current when challenged with 5-HT^+^ or cocaine. However, unlike in D437T and S438A, the amplitudes of the 5-HT^+^- and cocaine-induced upward deflection were equal. (**E**) Representative traces of currents through D437T in the presence of 150 mM K^+^_in_ recorded in the voltage range between –80 mV and 10 mV with increments of 10 mV. At each voltage 30 µM cocaine was applied. Traces were baselined to the current level in the presence of cocaine. (**F**) Current-voltage plot of the normalized cocaine-induced upward deflection of the current measured from D437T (*black circles*), S438A (*red squares*), and T284A (*teal triangles*), as a function of the membrane potential in the presence of 150 mM K^+^_in_ (n = 13, n = 12, n = 5, respectively). The increase in current amplitude at more negative voltages is consistent with the presence of a substrate-independent constitutive current.

In wild-type SERT, the transient peak was followed by the expected, K^+^-dependent, steady inward current (**Fig. 1A**). In contrast, in all three mutants, continued substrate application produced an upward deflection of the current that persisted for the duration of substrate exposure. To determine whether this upward deflection reflected inhibition of a constitutive conductance, we applied a saturating concentration of cocaine (30 µM) following substrate addition. Cocaine produced a similar upward deflection in cells expressing D437T, S438A or T284A (**Figs. 7B-D**), but not in cells expressing wild-type SERT (**Fig. 7A**). The most parsimonious interpretation is that all three mutants support a substrate-independent constitutive current that is blocked by the inhibitor. Consistent with this interpretation, the magnitude of the current through each mutant increased at more negative potentials (**Fig. 7E**).

In mutants that retained a transport-associated peak current (D437T and S438A), inhibition by 5-HT^+^ was smaller than that produced by cocaine, indicating that substrate only partially blocks the constitutive current. By contrast, in T284A, inhibition by 5-HT^+^ closely matched that elicited by cocaine, suggesting a more complete block of the constitutive conductance.

### The constitutive current is carried by Na^+^ ions

Because the Ct2-site appears to form a constriction between the extracellular and cytoplasmic sides of the transporter in some conformational states (**Fig. 4F-J**), we considered whether Na^+^ ions might permeate through SERT to give rise to the constitutive currents observed in all three mutants. We therefore recorded currents through D437T, S438A, and T284A at increasing extracellular Na^+^ concentrations. Representative recordings (**Fig. 8A-C**) show that the constitutive current was not detectable in the absence of extracellular Na^+^ and increased progressively in magnitude as the Na^+^ concentration was raised, approaching saturation at concentrations above 150 mM. The Na^+^ dependence of the normalized amplitudes of the constitutive current (**Fig. 8D**) was well described by a Hill equation. Fits yielded *K*_M_ values of ∼60 mM and Hill coefficients of >2 (see **Fig. 8** legend). Together, these measurements indicate that the constitutive current observed in these mutants is carried by Na^+^ ions and is consistent with permeation involving more than one Na^+^ ion.

**Figure 8.**
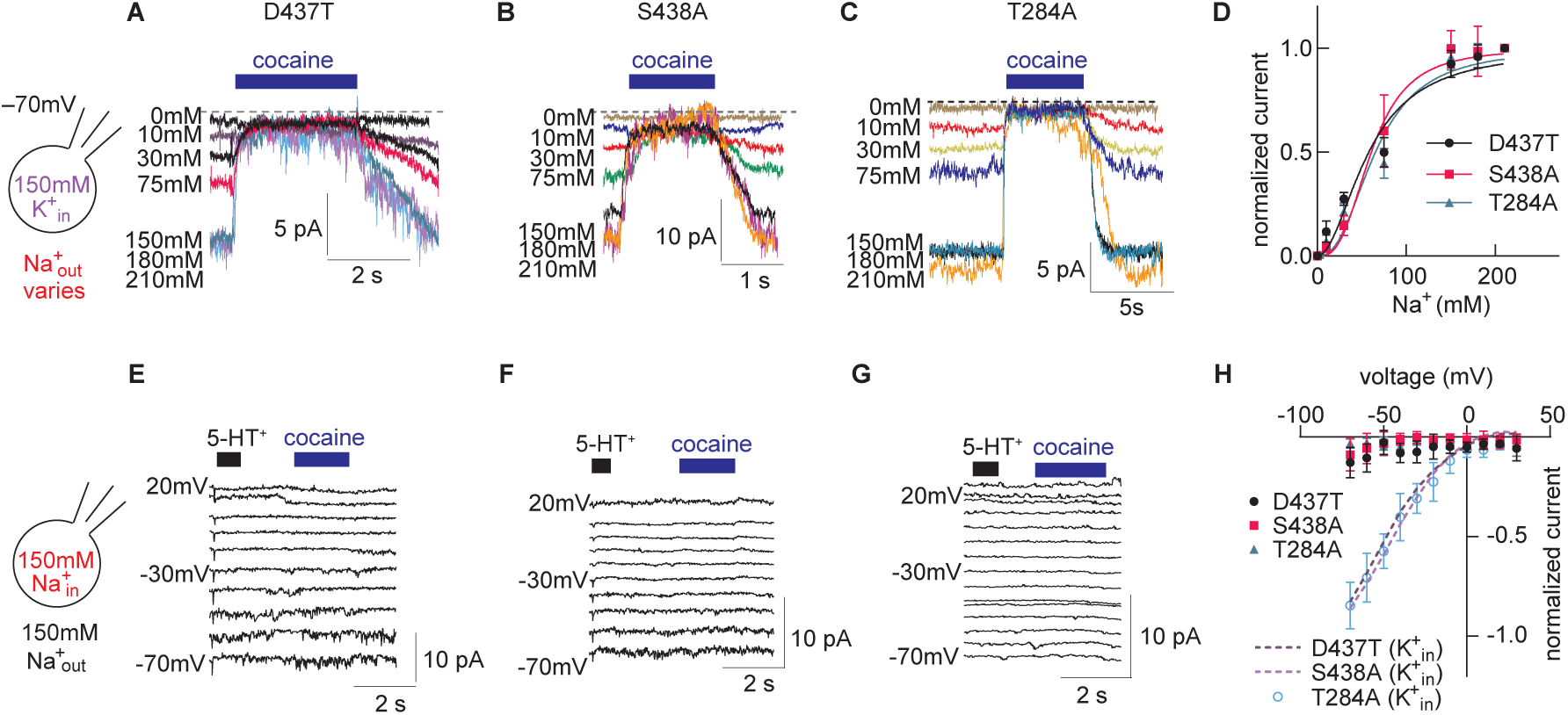
The constitutive, uncoupled current through the Ct2 pathway mutants is carried by Na^+^ ions. (**A-C**) Representative recordings obtained from a cell expressing D437T (A), S438A (B) or T284A (C). The substrate-independent constitutive current through each mutant was defined by the application of 30 µM cocaine (*blue bar*) and measured at 0, 10, 30, 75, 150, 180, and 210 mM Na^+^_out_, respectively. Traces were baseline-corrected (*dashed line*) to the current level in the presence of cocaine. The schematic shows the experimental conditions used in these experiments (*left*). (**D**) Normalized constitutive currents obtained from experiments such as (A-C) as a function of the Na^+^_out_ concentration. Data were fitted to the Hill equation (*black line*). The K_M_ values and the Hill slopes estimated by the fits were 57.8 ± 4.1 mM and 1.96 ± 0.18 for D437T (n = 5), 59.0 ± 4.0 mM and 2.20 ± 0.07 for S438A (n = 7), and 65.3 ± 6.2 mM and 2.20 ± 0.13 for T284A (n = 9). (**E, F, G**) Representative recordings obtained from cells expressing D437T (E), S438A (F), and T284A (G) in the presence of 150 mM Na^+^_in_ using the experimental conditions shown in the schematic (*left*). Currents were assessed at potentials ranging from –70 mV to 20 mV. Addition of 5-HT^+^ (*black bar*) evoked the peak current in D437T and S438A, but not in T284A. 30 µM of cocaine (*blue bar)* did not produce a visible change in the current amplitude, showing that, in the presence of 150 Na^+^_in_, the constitutive current was absent in all mutants. (**H**) Current-voltage plot for the voltage dependence of the constitutive current in the presence of 150 mM Na^+^_in_ (*filled symbols*: black *circles* D437T (n = 5), *squares* are S438A (n = 6), *triangles* are T284A (n = 5), compared to the presence of intracellular K^+^ (*dashed lines and open circles*) shown as smoothened data of that in Fig. 7F.

If the constitutive current is carried by Na^+^ ions, it should be eliminated when the Na^+^ driving force is removed. We therefore increased the intracellular Na^+^ concentration to 150 mM to match the extracellular Na^+^ concentration and thereby abolish the transmembrane Na^+^ gradient. Under these conditions, rapid application of a saturating concentration of 5-HT^+^ (30 μM) still elicited a transport-associated peak current in D437T (**Fig. 8E**) and S438A (**Fig. 8F**), but not in T284A (**Fig. 8G**), while abolishing the constitutive current in all three mutants. Moreover, superfusion with cocaine produced no change in current over the voltage range examined (–70 mV to +20 mV; **Fig. 8H**). Together, these results demonstrate that the constitutive current observed in the Ct2-pathway mutants is driven by an inwardly directed Na^+^ flux that depends on the transmembrane Na⁺ gradient.

### The constitutive current through the Ct2-pathway mutants represents Na^+^ uniport

A constitutive current can arise either from ions permeating through a channel-like state of the transporter, as observed for Li^+^-induced currents in wild-type SERT (Mager et al. 1994), or from futile transport cycles in which Na^+^ is transported in the absence of substrate, i.e., uniport or slippage. A defining feature of channel-like behavior is reversal of current direction with voltage. Consistent with this, Li^+^-induced currents in wild-type SERT reversed at positive potentials (**Fig. 9**).

**Figure 9.**
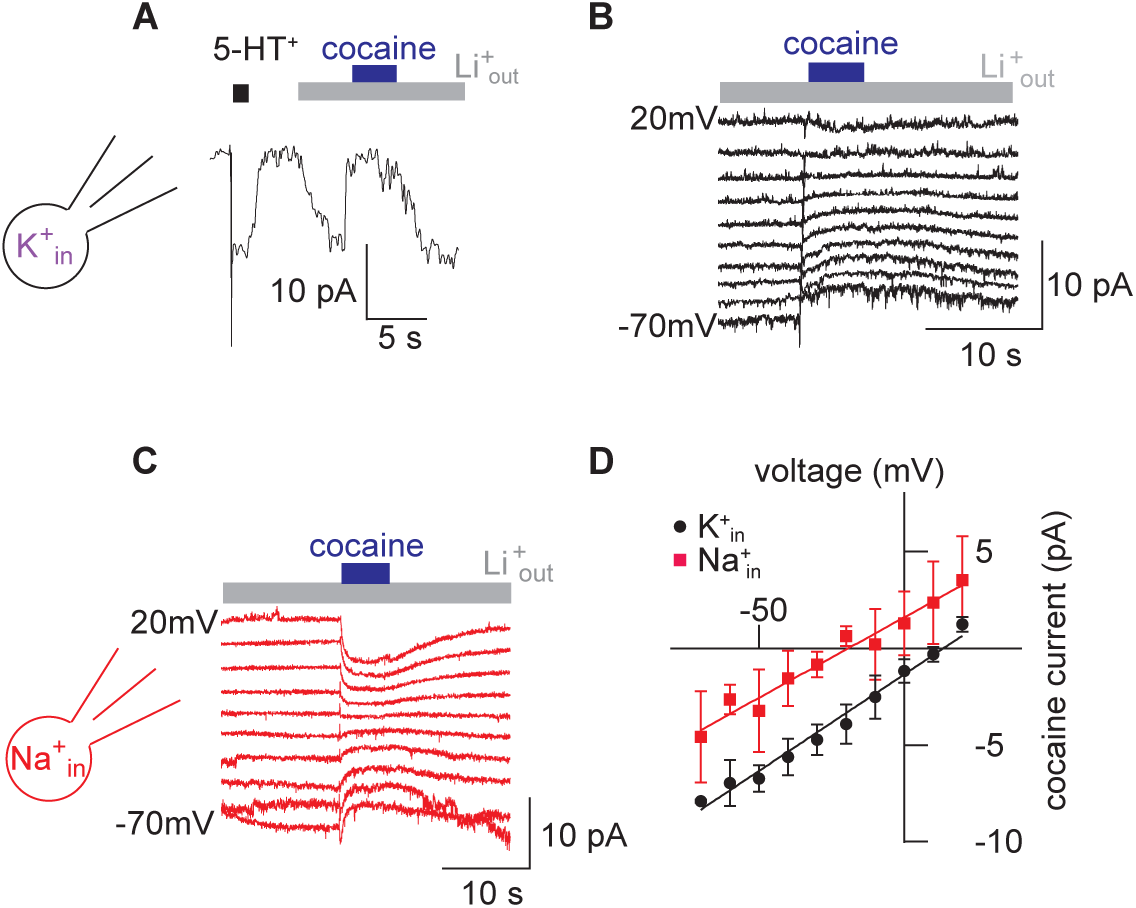
The channel-like behavior of the Li^+^-induced constitutive current through wild-type SERT. (**A**) Representative current recorded from a cell expressing wild-type SERT at –70 mV. The electrode solution contained 150 mM K^+^ (see schematic, *left*). In the presence of 150 mM Na^+^_out_, addition of 30 µM of 5-HT^+^ (*black bar*) resulted in an inwardly directed peak and steady current characteristic of wild-type SERT. Replacement of Na^+^_out_ with 150 mM Li^+^ (*gray bar*) gave rise to an inwardly directed constitutive current. Concurrent addition of 30 µM of cocaine blocked the constitutive current and subsequent removal of cocaine resulted in the reappearance of the constitutive current. (**B**) The Li^+^ induced current through wild-type SERT recorded at potentials ranging from –70 mV to 20 mV under the same conditions as in (A). (**C**) The Li^+^ induced current through wild-type SERT in the presence of 150 mM Na^+^_in_. (**D**) Current-voltage relationship of the Li^+^ induced current in the presence of 150 mM K^+^_in_ (*black circles*) and 150 mM Na^+^_in_ (*red squares*). Lines are linear fits to the data points. The slope and *y*-intercept of the black and red lines are 0.083 ± 0.006 pA/mV and –19.25 ± 3.44 mV, and 0.100 ± 0.006 pA/mV and 13.48 ± 2.68 mV, respectively.

By contrast, the constitutive currents elicited by D437T, S438A and T284A did not reverse direction over the voltage range examined (**Fig. 7F**) and instead exhibited pronounced inward rectification. Moreover, increasing the intracellular Na^+^ concentration to 150 mM, a condition anticipated to saturate any inward-facing Na^+^ sites, shifted the current-voltage relationship of the Li^+^-induced current to more negative potentials (**Fig. 9D**), whereas it reduced the amplitude of the mutant currents across a broad voltage range (**Fig. 8H**). These distinct responses indicate that the Li^+^-induced current and the constitutive current arising from mutations in and around the Ct2 site reflect different underlying mechanisms.

Importantly, the reduction of current amplitude at high intracellular Na⁺ concentrations in the mutants is consistent with Na^+^ uniport. In this scenario, elevated occupancy of inward-facing Na^+^ sites is expected to impede completion of the transport cycle and thereby slow Na^+^ influx, leading to reduced current amplitudes. This interpretation is further supported by the reduced concentrative power of the mutants (**Fig. 2C-E**), consistent with uncoupled Na^+^ transport.

### Disrupting the TM5-TM8 connection has a distinct effect on transport

Although uptake was measurable for all three mutants, T284A exhibited the lowest concentrative power (**Fig. 2C**). This phenotype is consistent with the absence of substrate- and Na^+^-dependent peak currents in T284A at the resolution of the patch clamp recordings (**Fig. 8C**). In addition, the ability of 5-HT^+^ to fully block the constitutive current was unique to T284A (**Fig. 7B-D**). Together, these observations indicate that the disruption of the Thr284-mediated TM5–TM8 interaction has consequences that are distinct from perturbations of the Ct2 site residues Asp437 and Ser438.

To further probe these differences, we examined the temperature dependence of the substrate-free currents by recording conductance at different temperatures. In wild-type SERT, both peak and steady-state currents increased with temperature, and no constitutive currents are measurable even at 37°C (**Fig. 10A**). By contrast, all three mutants exhibited a temperature dependent increase in cocaine-sensitive conductance (**Fig. 10B-D**). Analysis of the Arrhenius relationships revealed that the apparent activation energy (E_A_) for the uniport-like event was higher for D437T and S438A (>30 kJ/mol) than for T284A (∼12 kJ/mol) (**Fig. 10F-H**).

**Figure 10.**
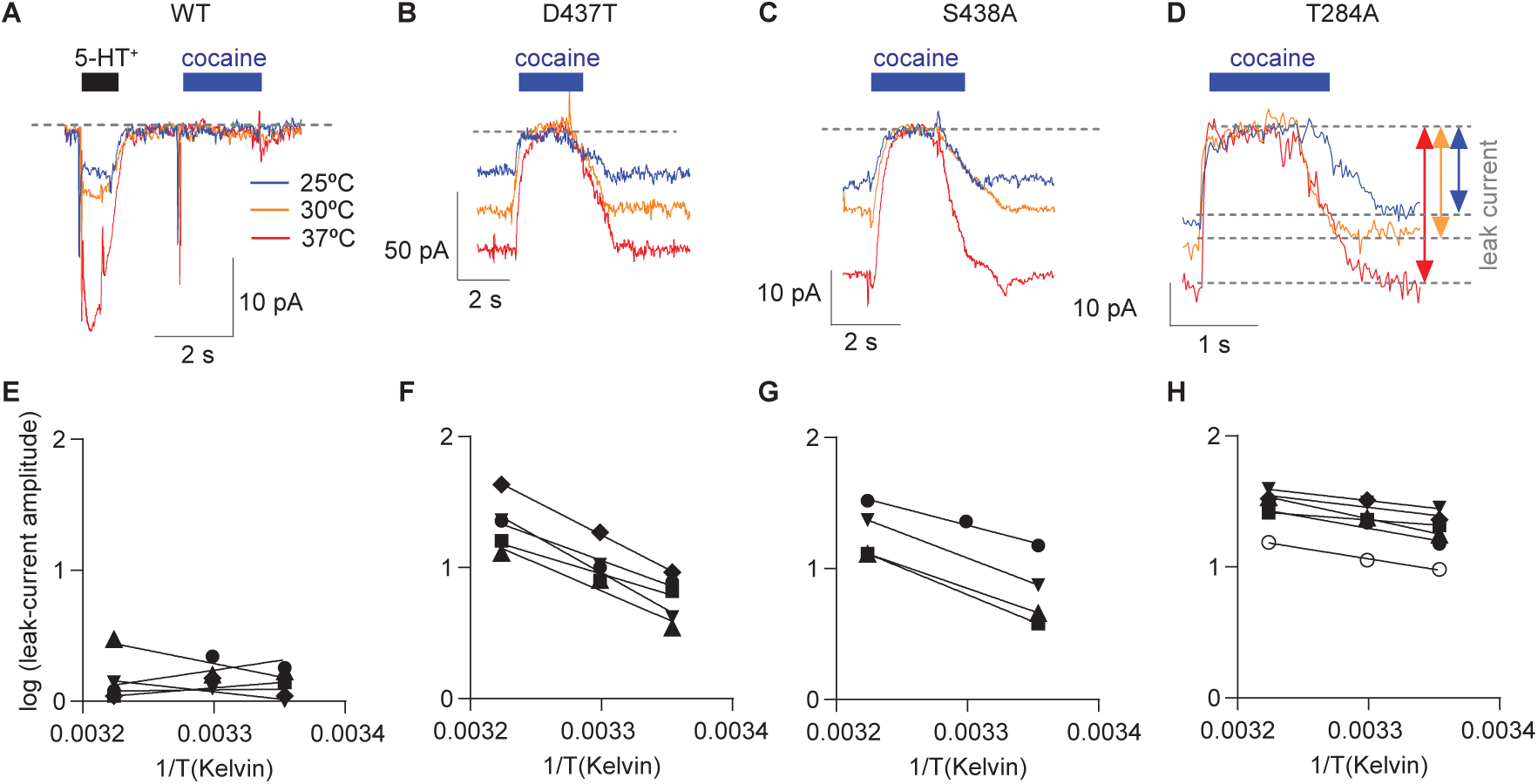
The activation energy of the substrate-independent constitutive current is higher in D437T and S438A than in T284A. (**A-D**) Recordings of the substrate-independent constitutive current through wild-type SERT (A), D437T (B), S438A (C), and T284A (D) at different temperatures (25°C, 30°C, and 37°C in *blue, orange,* and *red*, respectively). No constitutive current was detected in cells expressing wild-type SERT, so 30 µM of 5-HT^+^ was applied to wild-type SERT as a control for protein expression. (**E-H**) Arrhenius plots of the constitutive current amplitudes of SERT wild type (E), D437T (F), S438A (G), and T284A (H). Each line in a plot connects data points obtained from one independent experiment. The average activation energies (*E*_A_) were: 36 ± 8.7 kJ/mol for D437T (n = 5); 30 ± 5.4 kJ/mol for S438A (n = 4); and 11.9 ± 4.3 kJ/mol for T284A (n = 6). Panel E is shown for completeness; no constitutive current was detectable in wild type SERT.

These results indicate that breaking the TM5-TM8 connection via Thr284 lowers the energetic barrier for uncoupled Na^+^ flux to a greater extent than perturbations at the Ct2 site. By comparison, the absence of detectable constitutive currents in wild-type SERT suggests that the corresponding energetic barrier exceeds what can be overcome by thermal fluctuations, even at physiological temperatures.

## Discussion

### Ct2 links Na⁺ occupancy to local interaction networks

Although the thermodynamic role of Na⁺ in driving serotonin uptake has long been established, how occupancy of individual ion-binding sites controls SERT conformation remains incompletely understood. Previous biophysical and biochemical studies have shown that Na⁺, K⁺, and inert cations shift the populations of SERT conformations (Fenollar-Ferrer et al. 2014; Möller et al. 2019). Our combined mutational, electrophysiological, and simulation results identify local interaction networks through which Na⁺ occupancy at Ct2 may contribute to these shifts and maintain coupling between Na⁺ and 5-HT⁺ transport.

The simulations support two complementary networks (**Fig. S6**). Na⁺ occupancy favors a Ser438 orientation that permits hydrogen bonding to Tyr95 in TM1 and shapes the region between Ct2 and S1. Na⁺ also restricts the orientation of Asp437, favoring its interaction with Thr284 in TM5. The involvement of Thr284 is consistent with observations in other LeuT-fold transporters: residues in TM5 interact with the position equivalent to Asp437 in MhsT and BetP, supporting a link between cation binding and TM5 reorientation (Stolzenberg et al. 2017; Perez et al. 2014). The Thr284-Asp437 distance also emerged as a kinetically relevant feature in adaptive sampling simulations of 5-HT^+^ import by SERT (Chan et al. 2022).

Together our results support a working model in which these networks coordinate access to Ct2, helping prevent Na^+^ leakage before 5-HT^+^ binding (**Fig. 11A-C**). In the substrate-free outward-open state, Ct2-bound Na^+^ would favor the Ser438-Tyr95 interaction, restricting Na^+^ escape toward the extracellular solution. At the same time, the Asp437-Thr284 interaction would stabilize the helical structure of TM5 and maintain closure of the cytoplasmic pathway.

**Figure 11:**
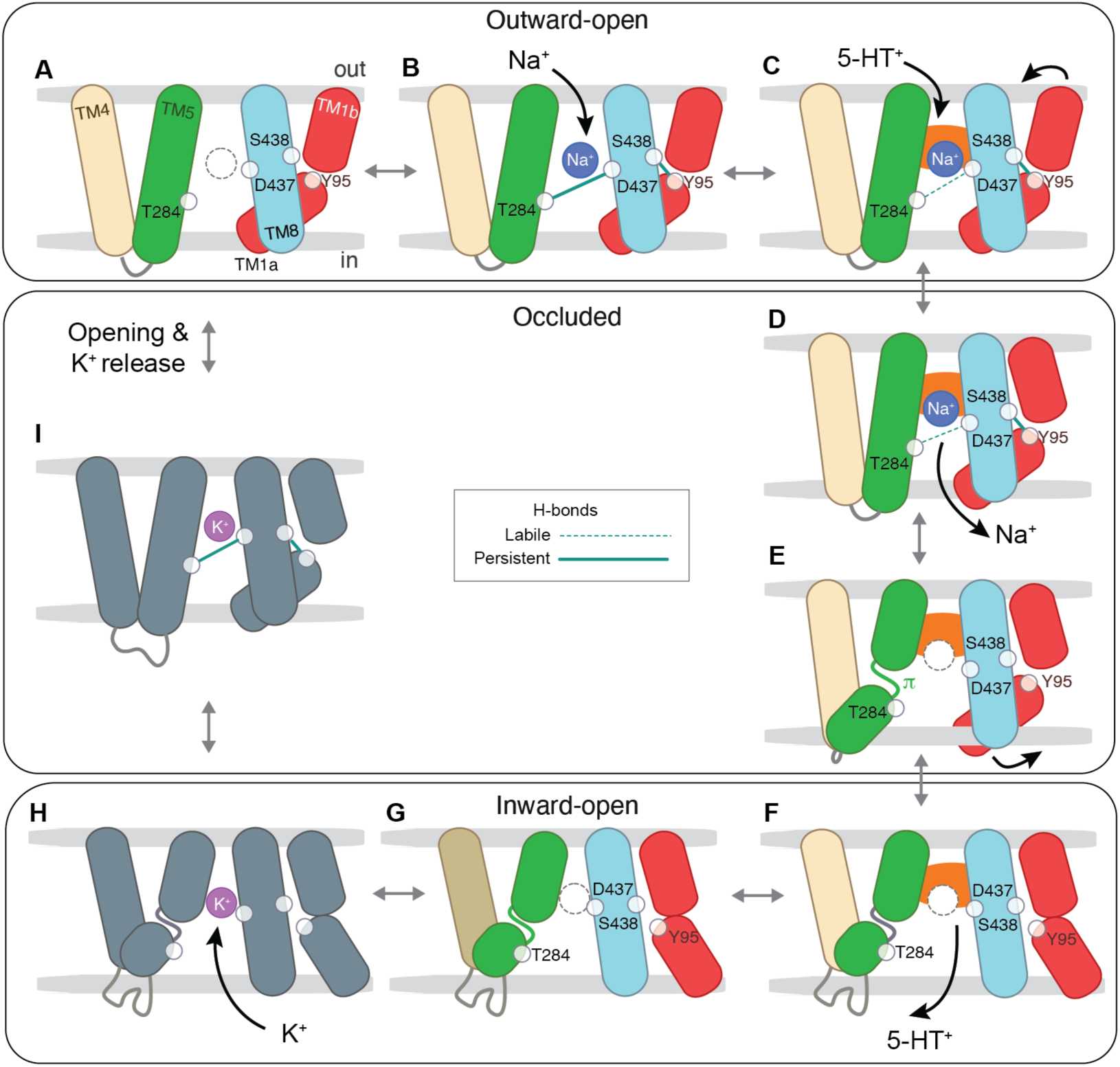
Working model of the interactions proposed to connect Ct2-site binding to distal conformational changes during the SERT transport cycle. Transmembrane helices TM1a and TM1b (*red*), TM4 (*wheat*), TM5 (*green*), and TM8 (*blue*) are shown as ovals. States not simulated are shown using gray ovals. Membrane boundaries are indicated with gray horizontal bars. H-bonds between residues Thr284-Asp437 and Ser438-Tyr95 (*small white circles*) are indicated as solid teal lines where persistent or dashed teal lines where labile (infrequent or exchanging); no line indicates the bond is not formed. **(A-C)** Outward-open conformations with Ct2 empty and no 5-HT^+^ (A; *dashed circles indicate empty sites*), with Na^+^ bound at Ct2 (B; *blue circle*) and with both Na^+^ and 5-HT^+^ (C; *orange*) bound. Panel A should be interpreted cautiously as it is difficult state to capture (*n* = 1). **(D-E)** Outward-occluded conformations with Na^+^ and 5-HT^+^ bound (D) and after Na^+^ release into the cytoplasm (E); π marks the π-bulge in TM5. **(F-H)** Inward-open conformations with 5-HT^+^ bound (F), after 5-HT^+^ unbinding (G), and with K^+^ bound (H; *purple circle*). **(I)** K^+^-bound occluded “return” state, which returns to the outward-open (A) on opening of the extracellular pathway and release of K^+^. In (I), indicated H-bonding is inferred from the equivalent Na^+^-bound state. Double-headed arrows indicate proposed reversible transitions; the simulations do not establish their order. Panels A and E are likely short-lived states.

Following 5-HT^+^ binding and extracellular pathway closure, local unwinding and separation of TM5 from TM8 could facilitate hydration of a cytoplasmic route for Na^+^ release (**Fig. 11D-G**). The requirement for unwinding at TM5 is supported by the deleterious effects on transport activity and conformational equilibria of mutations at both Pro288 and Gly278 (Zhang et al. 2025), both of which are expected to confer conformational flexibility to TM5. Na^+^ loss would then further weaken the Asp437-Thr284 interaction and disfavor the Ser438-Tyr95 interaction, helping release the restraints on TM5 and TM1a required for cytoplasmic opening and substrate release. The simulations support the local interactions underlying this model but do not directly establish the sequence of transitions.

### Mutant currents suggest premature access to Ct2 from the cytoplasm

All three mutants exhibit constitutive currents, indicating that Na^+^ movement can occur without 5-HT^+^ transport. Why then does Na^+^ not conduct through a continuously open channel, as proposed for Li^+^? The mutants retain some ability to concentrate 5-HT^+^ (**Fig. 2**) and both 5-HT^+^ and cocaine inhibit their constitutive currents (**Fig. 7**). These observations suggest that the constitutive current remains sensitive to ligand occupancy and that aspects of the coupled transport cycle are preserved. Together with the simulation results, they support a model of repeated Na⁺ binding and release rather than a continuously open pathway. Indeed, the remaining Ct2 coordinating groups appear to still support Na⁺ binding and conformational cycling: D437T retains a hydroxyl-bearing side chain at this position, as found in other Na⁺-coupled SLC6 transporters, whereas S438A retains Asp437. Nevertheless, although altered affinity may be a factor, the combined results support a specific role for Asp437, Ser438 and Thr284 in converting Na⁺ occupancy into the pathway closure needed to prevent leakage before substrate capture.

We propose that the mutants weaken the interactions that normally prevent Na^+^ leakage from substrate-free, outward-open SERT. In D437T, reduced interactions with Thr284 could allow TM5 to separate more frequently from TM8, permitting a Ct2-bound Na^+^ ion to escape to the cytoplasm before 5-HT^+^ binding and extracellular pathway occlusion. In S438A, loss of the Ser438-Tyr95 interaction could similarly favor local separation at these positions and facilitate cytoplasmic access.

If extracellular access remains available after Na^+^ escapes, another Na^+^ ion could occupy the Ct2 site and restore closure of the cytoplasmic pathway. Repeated binding and release could therefore permit Na^+^ uniport without requiring a continuously open channel. The present data do not distinguish simple Ct2 reoccupation from mechanisms involving additional Na⁺ occupancy near S1 or a knock-on effect.

### Ligand inhibition distinguishes the mutant phenotypes

Cocaine stabilizes outward-open SERT and occupies the S1 pocket (Yang et al. 2023). Its complete inhibition of the mutant currents is consistent with blockade of the extracellular pathway to the Ct2 site, interrupting the repeated Na^+^ access required to sustain the constitutive current.

The incomplete inhibition by 5-HT^+^ in S438A and D437T suggests a different outcome. Unlike cocaine, 5-HT^+^ promotes progression toward occluded and inward-open conformations. We propose that these mutants permit premature Na^+^ release while extracellular access remains available during progression toward occlusion. Another possibility, especially in the case of S438A, is that Na^+^ can reach the Ct2 site by moving past 5-HT^+^ which occupies less of the large S1 pocket than cocaine.

By contrast, 5-HT^+^ completely blocked the constitutive conductance of T284A, indicating that substrate binding suppresses Na^+^ leakage more effectively in this mutant than in S438A or D437T. One possibility is that Na^+^ binding, while diminished, remains sufficiently robust in this mutant that the Na^+^ ion cannot unbind and move into the cytoplasm even upon formation of a 5-HT^+^-dependent occluded state. In the absence of 5-HT^+^, the loss of H-bonding potential in T284A increases the likelihood of TM5 separating from Asp437 in TM8, allowing the pathway to open more frequently and explaining the lower energetic barrier for uniport by this mutant. Together with the simulation results, this phenotype supports a role for Thr284 in coordinating the position of TM5 with Na^+^ occupancy to prevent leakage before substrate capture in SERT.

### Simulation scope and unresolved ion stoichiometry

Several features of native SERT were omitted from the simulations. The input structures do not resolve the segment N-terminal to residue 74, likely reflecting its considerable flexibility (Fenollar-Ferrer et al. 2014). The simulations also omitted cholesterol, additional extracellular substrates and plasma-membrane like lipid mixtures. Cholesterol is commonly observed in structures with a closed cytoplasmic pathway, and a second serotonin molecule or detergent molecule has been detected in an extracellular region (Yang and Gouaux 2021; Yang et al. 2023). These simplifications reduced the sampling demands but limit the contexts in which the proposed mechanism has been tested. The omitted components may alter conformational populations (Rudin et al. 2025; Scanlon, Williams, and Schloss 2001); whether they also modify the local interaction networks described here remains to be established. Future work should examine these effects, including those of the N-terminal region, as investigated for DAT (Xu and Chen 2022; Khelashvili et al. 2015).

The ion stoichiometry of SERT also remains unresolved. The recent proposal that Cl^−^ acts as a co-factor rather than a transported ion(Huang et al. 2026) raises a charge-balance question: inward transport of one Na^+^ ion and one 5-HT^+^ molecule coupled to outward transport of one K^+^ ion would yield net inward positive charge, whereas the functional transport cycle has been reported to be electroneutral (Rudnick and Nelson 1978). In a recent study using simulations of substrate-free SERT, protonation of Asp98 stabilized the ions in the K^+^-bound state (Hellsberg et al. 2024). We therefore considered whether outward H^+^ movement could provide the missing charge compensation. However, measurements of APP^+^ uptake into SERT proteoliposomes did not exhibit the cumulative effects of H^+^ and K^+^ gradients expected under this proposal (Hellsberg et al. 2024). Other studies instead place K^+^ at Ct1 (Schmidt et al. 2024) or support Cl^−^ cotransport (Tan et al. 2026). Resolving these alternatives will be necessary to place the Ct2 mechanism within a complete transport cycle.

In conclusion, our biochemical, electrophysiological and simulation results support a molecular model in which Ct2-dependent interaction networks coordinate structural changes with Na^+^ occupancy. Disrupting these networks permits Na^+^ flux without substrate transport, identifying a mechanism by which SERT normally confers efficient Na^+^-5-HT^+^ coupling as expected for healthy synaptic function.

## AI statement

During the preparation of this work, the authors used Claude and ChatGPT to suggest clearer language in the introduction, results and discussion. The authors reviewed and edited the output as needed and take full responsibility for the content of the published article.

## Acknowledgments

We thank Gary Rudnick for useful discussions. This research was supported in part by the Austrian Science Foundation Grant 10.55776/P36667 and the Intramural Research Program (ZIA NS003139) of the National Institutes of Health (NIH). The contributions of the NIH author(s) were made as part of their official duties as NIH federal employees, are in compliance with agency policy requirements, and are considered Works of the United States Government. However, the findings and conclusions presented in this paper are those of the author(s) and do not necessarily reflect the views of the NIH or the U.S. Department of Health and Human Services. This work utilized the computational resources of the NIH HPC Biowulf cluster.

## Funding

National Institutes of Health, National Institute of Neurological Disorders and Stroke,

Division of Intramural Research NS003139 (LRF)

Austrian Science Foundation doi:10.55776/P36667 (WS)

## Author contributions

Conceptualization: WS, EH, DB, LRF

Molecular simulations: EH, AB, JH, JS, NB, LRF

Simulation analysis: EH, AB, JH, JS, NB, LRF

Bench experiments: DB, EE

Visualization: EH, AB, JS, LRF

Supervision: EH, WS, LRF

Writing—original draft: EH, WS, LRF

Writing—review & editing: all authors

Resources, project administration and funding acquisition: WS, LRF

## Competing interests

The authors declare that they have no competing interests.

## Notes

### Competing Interest Statement

The authors have declared no competing interest.

