## Supplementary Materials for "Sodium coordination at the Na2 site enforces ion-substrate coupling in serotonin transporter SERT"

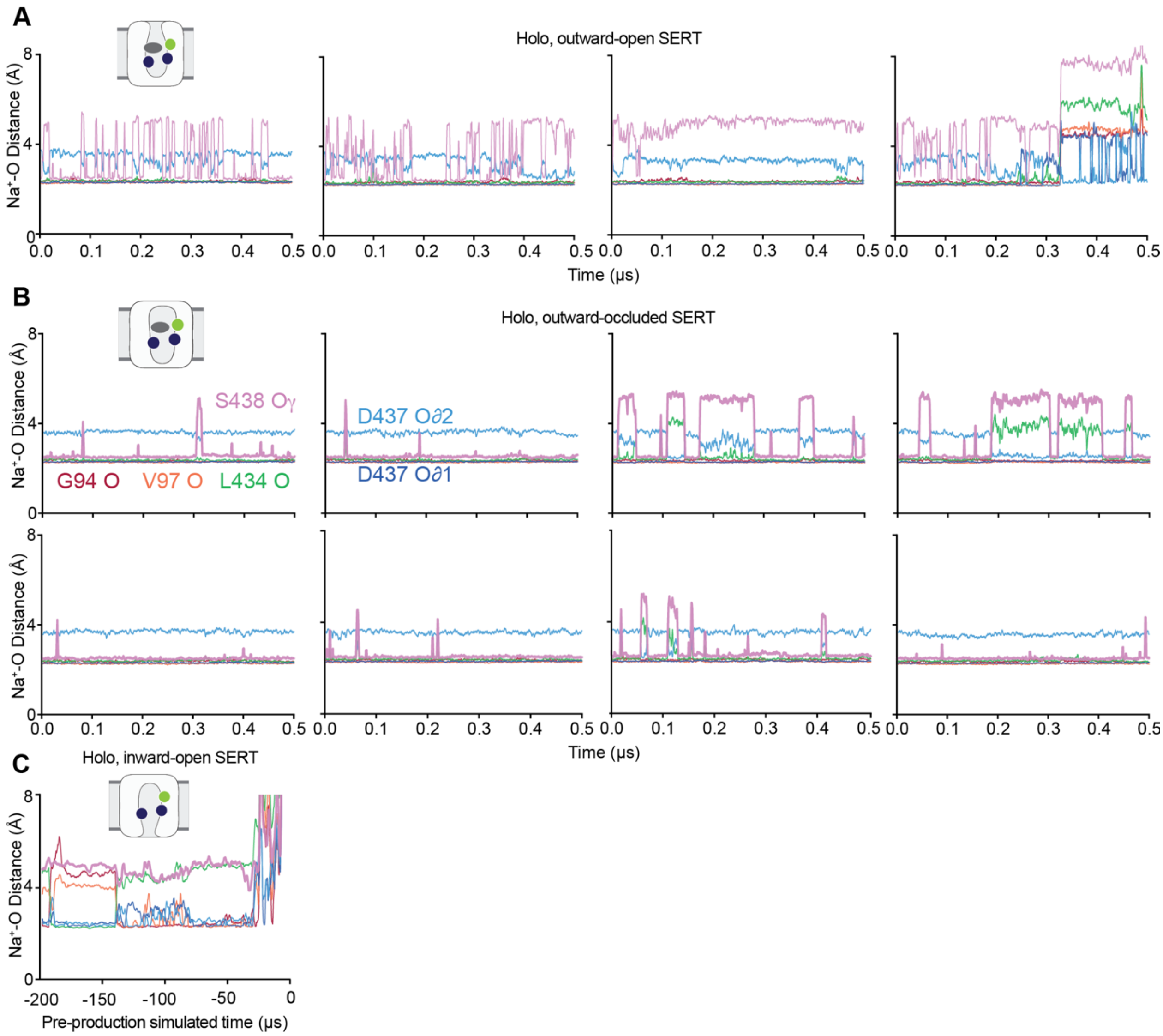


**Fig. S1. Time dependence of Ct2 site $Na^+$-O distances during MD simulations of SERT bound to 5-$HT^+$, two $Na^+$ ions, and a $Cl^-$ ion.**

The transporter was in an outward-open **(A),** outward-occluded **(B)** or inward-open **(C)** conformation. Traces indicate distances from the backbone O atoms of Gly94 (*red*), Val97 (*orange*), and Leu434 (*green*) and the side chain O atoms of Asp437 (*blues*) and Ser438 (*violet*). (A, B) Production replicas ($n = 4$; A, $n = 8$; B). (C) Pre-production equilibration, with progressively released restraints on the ion (see Methods).

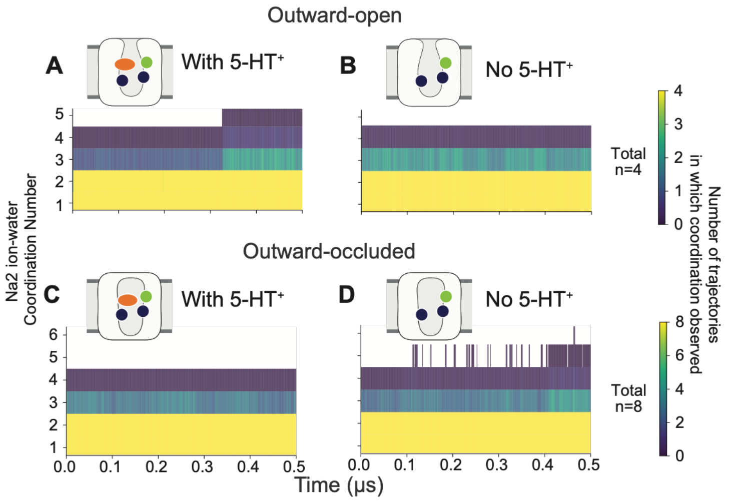


**Fig. S2. Number of water molecules coordinating the $Na^+$ ion at the Ct2 site during MD simulations of SERT.**

The transporter was in (**A, B**) outward-open or (**C-D**) outward-occluded conformations as indicated in the schematics. The number of trajectories (relative to the total indicated in the legend) in which a given number of waters is observed, is plotted as a function of simulation time. For example, the yellow region from $y = 1, 2$ signifies that two water molecules coordinate the ion in all frames in all trajectories, whereas higher water counts are less common. Coordination is defined as a water O within 2.3 Å of the Ct2 $Na^+$ ion. The S1 substrate site is occupied either with 5-$HT^+$ (A,C; *orange oval in schematic*) or no ligand (Asp98 deprotonated; B,D).

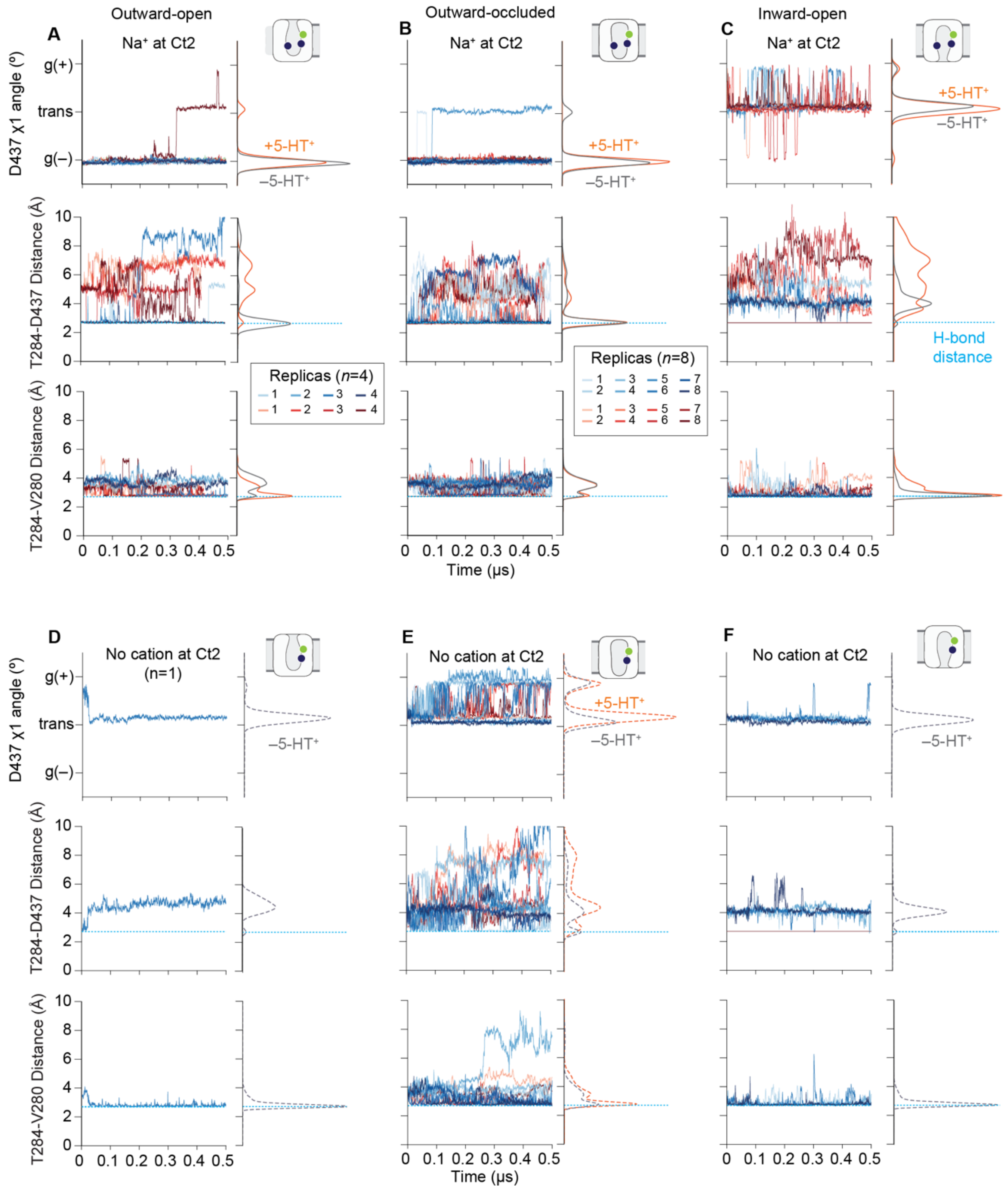


**Fig. S3 Time-resolved behavior of the Ct2–TM5 interaction network across conformational states.** Time series of the Asp437 χ1 dihedral angle (*top row of each panel*), Thr284–Asp437 distance (*middle row*), and Val280–Thr284 distance (*bottom row*) from individual MD trajectories. Corresponding population distributions are shown to the right of each plot and reproduce the data shown in **Fig. 3**. **(A–C)** Simulations with $Na^+$ at the Ct2 site in the outward-open (A), outward-occluded (B), and inward-open (C) conformations, as indicated by the schematics. **(D–F)** As in (A–C) but for simulations with no cation at Ct2. Coloring indicates the S1-site condition: Apo (*blue*) or 5-$HT^+$ bound (*orange*). Each trajectory is 500 ns and $n = 4$ or $n = 8$. Horizontal dashed lines in distance panels indicate H-bonding distance (2.7 Å).

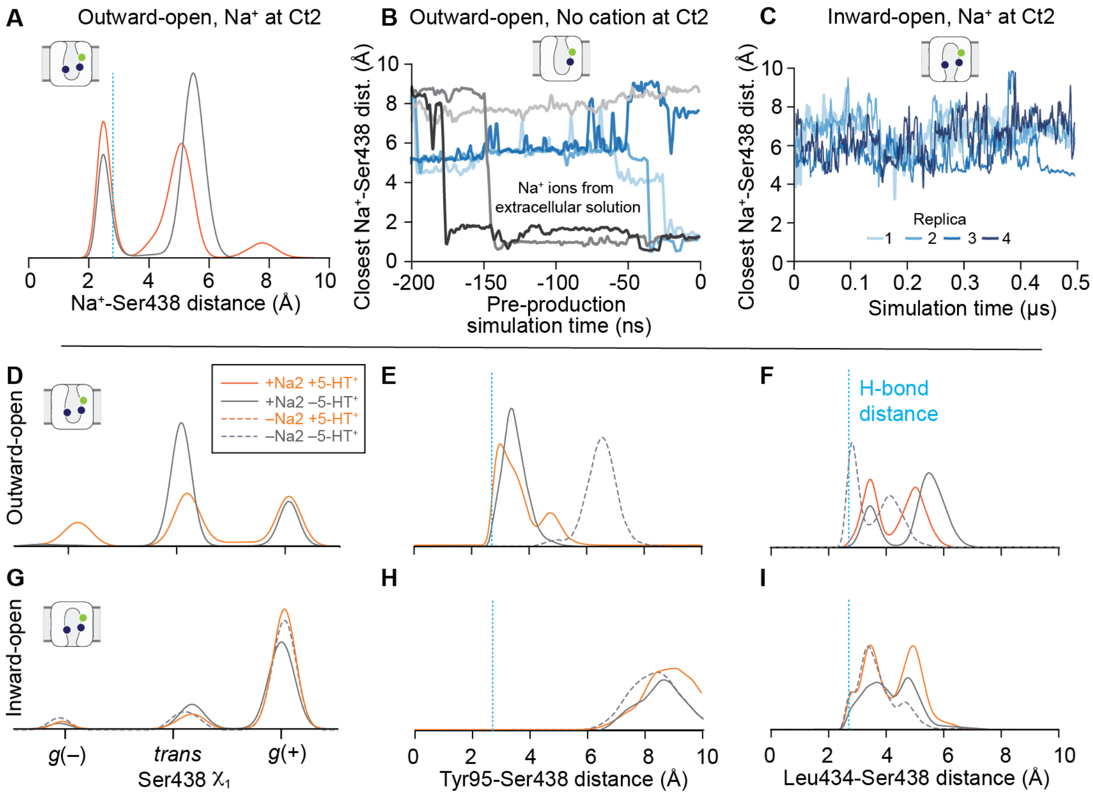


**Fig. S4. Characterization of Ser438 dynamics during MD simulations of SERT.**
**(A-C)** Distance from the side chain hydroxyl O atom of Ser438 either to the $Na^+$ ion initially positioned at Ct2 or to the closest cation at any frame in the trajectory, for simulations of the outward-open (A-B) and inward-open (C) states. (**A**) Distribution of distances for the outward-open state measured in the presence (*orange*) or absence (*gray*) of 5-$HT^+$. Underlying time traces for the 5-$HT^+$-bound system are shown in **Fig. S1**. Equivalent ion-hydroxyl distances for the outward-occluded state are shown in **Fig. 2B**. **(B)** Spontaneous entry of cations from the aqueous solution into the substrate pocket during equilibration of the outward-open state without 5-$HT^+$ or a $Na^+$ at Ct2 when no restraint was imposed (**see Methods**). **(C)** In the inward-open state, $Na^+$ ions did not interact with the side-chain of Ser438. Similar data were observed in the presence of bound 5-$HT^+$. **(D-I)** Ser438 dihedral angle and distance distributions for (D-F) outward-open, or (G-I) inward-open states of SERT plotted as probability distributions where a $Na^+$ ion was placed at or near Ct2 (*solid lines*) or omitted (*dashed lines*). Equivalent data for the outward-occluded state are shown in **Fig. 6**. Simulations were carried out with the S1 site occupied by 5-$HT^+$ (*orange*) or no ligand (*gray*). **(D, G)** Distribution of side chain $\chi_1$ dihedral angle of Ser438. **(E-F, H-I)** Distance between the side chain oxygen of Ser438 and the backbone carbonyl oxygen of either Tyr95 in TM1a (E, H) or Leu434 in TM8 (F, I).

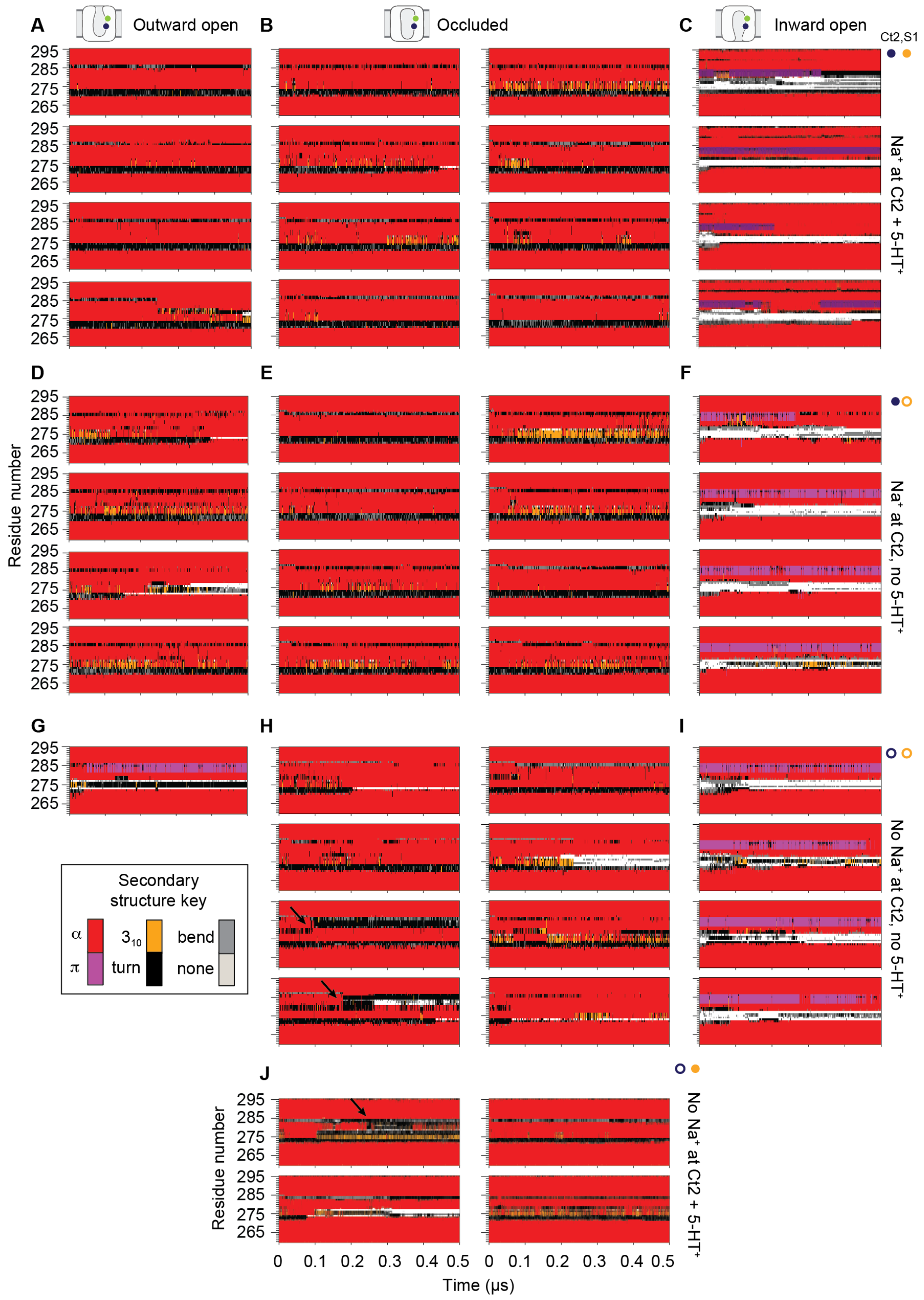


**Fig. S5. Time-resolved DSSP assignments for the TM4-5 loop region of SERT.** Time-dependent DSSP secondary-structure assignments are shown for individual MD trajectories of SERT in (**A, D, G**) outward-open, (**B, E, H, J**) outward-occluded, and (**C, F, I**) inward-open conformations. Each panel displays the temporal evolution over 0.5 µs of DSSP states (α-helix, $3_{10}$-helix, turn, bend, or no secondary structure) for residues 260-295, encompassing the turn region in TM4, the TM4-5 loop, and the proximal TM5 region. Each horizontal strip corresponds to one simulation replica (*n* = 1, 4 or 8 per condition), with colors indicating DSSP classifications as defined in the key. Four ligand/ion conditions were tested: the Ct2 site was occupied by a $Na^+$ ion (**A–F**) or left unoccupied (**G-J**). The S1 site contained 5-$HT^+$ (**A-C, J**) or no ligand (**D-I**). Arrows indicate timepoints where the proximal TM5 region converts to a turn. These time-resolved data underlie secondary-structure occupancies shown in **Fig. 4**.

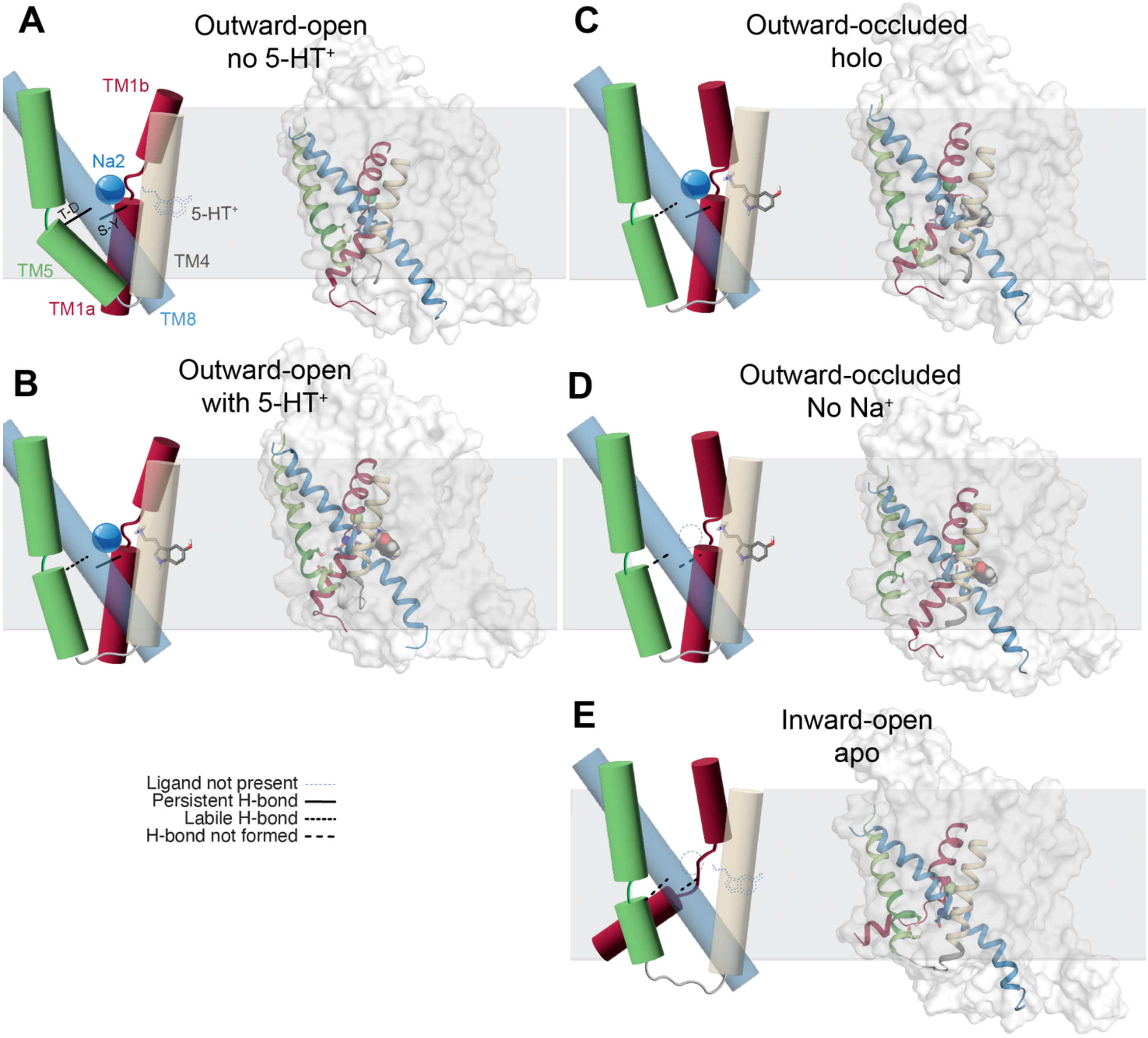


**Fig. S6. Overview of the conformational states involved in coupling Ct2 occupancy to TM5 and TM1a.**
Schematic and structural views of the **(A,B)** outward-open, (**C,D**) outward-occluded and (**E**) inward-open states. (A) and (B) represent the $Na^+$-bound outward-open state and the effect of binding 5-$HT^+$. (C) and (D) represent the 5-$HT^+$-bound occluded state and the effect of $Na^+$ loss. (E) Illustrates the opening of the main cytoplasmic pathway with TM1a displaced and the TM4-5 loop unstructured. In the structural view, selected helices are shown in the context of the overall protein surface; relevant side-chains are shown as sticks; serotonin and ions are shown using spheres when present. When absent, substrates and ions are indicated using thin dotted outlines in the schematics. For each conformation, the persistence (solid line), lability (dotted line), or absence (dashed line) of key H-bonds connecting TM8 with either TM5 or TM1a are indicated in the schematics: T-D refers to Thr284-Asp437 and S-Y refers to Ser348-Tyr95. Snapshots are taken from the last frame of one replica with each configuration.
